# MEK inhibition induces AXIN1 loss in colorectal cancer by mTOR associated suppression of protein synthesis

**DOI:** 10.1101/2025.06.13.659446

**Authors:** Nachiyappan Venkatachalam, Li Wang, Niyumi Muthukumarana, Robert Ihnatko, Jeroen Krijgsveld, Olga Skabkina, Antonia Leipertz, Yubin Chen, Johannes Betge, Kim Boonekamp, Michael Boutros, Georg Stoecklin, Matthias Ebert, Johanna Schott, Tianzuo Zhan

## Abstract

AXIN1 is a central regulatory hub of many oncogenic pathways in colorectal cancer (CRC). As the main scaffold protein and least abundant component of the beta-catenin destruction complex, changes in AXIN1 levels affect Wnt signaling output. We show that targeting the Ras-MAPK pathway by MEK1/2 inhibitors induces AXIN1 loss across a panel of CRC cell lines and patient-derived organoids. GSK3B inhibition similarly reduced AXIN1 levels, yet by distinct mechanisms. MEK1/2 causes a reduction of *AXIN1* transcript levels, but neither affects protein stability nor post-translational modifications of AXIN1. In contrast, GSK3B inhibition induces rapid AXIN1 degradation. Prevention of AXIN1 loss by co-treatment with tankyrase inhibitors was much stronger for GSK3B than for MEK1/2 inhibition. Using isogenic CRC cell lines and murine intestinal organoids, we show that APC truncations strongly reduce basal AXIN1 levels, but do not alter dynamics of AXIN1 loss upon MEK1/2 inhibition. Polysome profiling and Ribo-Seq revealed that MEK1/2 inhibition reduces global protein synthesis via an mTOR dependent pathway. This translational repression is sufficient to cause significant AXIN1 loss, as treatment with mTOR inhibitors phenocopies the effect of MEK1/2 inhibitors. Our study demonstrates that AXIN1 protein homeostasis is critically controlled by Ras-MAPK signaling at the level of protein synthesis, and that MEK1/2 inhibitors cause AXIN1 loss by translational repression.

## INTRODUCTION

Colorectal cancer (CRC) is a major cause of cancer-associated mortality (*1*). Approximately 20-30% of patients with CRC are diagnosed with metastatic disease and treated by pharmacotherapy (*2*). Currently, chemotherapy remains the main backbone of pharmacotherapy for most metastatic CRC (*3*). Despite the high frequency of Ras-MAPK pathway alterations in CRC, small molecules targeting this pathway such as BRAF or MEK1/2 inhibitors are less effective in CRC compared to other cancers harboring the same genetic alterations, e.g. melanoma or non-small cellular lung cancer (*4*, *5*). The underlying molecular mechanisms for the relative resistance of CRC to Ras-MAPK pathway inhibition are manifold, including genetic and non-genetic adaptation (*6*). We previously showed that MEK1/2 inhibitors induce stem cell plasticity by stimulating Wnt signaling in cell and organoid models of CRC, through a mechanism that involves reduction of cellular AXIN1 levels (*7*). AXIN1 is the main scaffold protein of the destruction complex, which consists of the core members APC, AXIN1/2, CK1α and GSK3B, and controls intracellular levels of beta-catenin by inducing proteasomal degradation (*8*). As the least abundant component of this multi-protein complex, AXIN1 levels critically determine the activity of canonical Wnt signaling. Decrease of AXIN1 levels was observed during ligand stimulation of the pathway (*9*) and functional depletion of AXIN1 levels increased canonical Wnt activity (*7*). Beyond Wnt signaling, AXIN1 also regulates other oncogenic pathways by controlling protein stability of key components (*10*). Together with GSK3B, PP2A and Pin1, AXIN1 stimulates ubiquitin-mediated degradation of the MYC oncogene (*11*). Similarly, AXIN1 acts with the ubiquitin-ligase RNF111/Arkadia to reduce levels of SMAD7, leading to activation of the TGF-beta pathway (*12*). Due to its role as a central regulatory hub for key oncogenic pathways, cellular levels of AXIN1 are regulated by diverse mechanisms. Transcriptional activation of the *AXIN1* gene is mediated by several transcription factors, such as GATA4 in osteoblasts (*13*), RUNX1 in breast cancer (*14*), PHB1 in CRC (*15*) and EGR1 in epithelial cell lines (*16*). Furthermore, protein homeostasis of AXIN1 is controlled by extensive post-translational modification and degradation via the ubiquitin-proteasome system (*10*). Most prominently, poly-ADP ribosylation (PARsylation) of AXIN1 by tankyrases and subsequent E3 ubiquitination by RNF146 was shown to mediate its degradation in CRC and other cancers (*17*). This process can be pharmacologically targeted by tankyrase inhibitors, resulting in a stabilization of AXIN1 and inhibition of Wnt signaling (*18*). Hence, AXIN1 is considered as a potentially druggable target of the Wnt pathway. Similarly, several other E3 ubiquitin ligases were discovered that target AXIN1 for proteasomal degradation, including SMURF2 (*19*), SIAH1/2 (*20*) and TRIM65 (*21*). Conversely, deubiquitination enzymes such as USP7 or USP34 stabilize AXIN1 and act as negative regulators of Wnt signaling (*22*, *23*). Given the critical, regulatory function of AXIN1 for Wnt signaling and its diverse interactions with other oncogenic pathways, it is important to mechanistically understand how pharmacological inhibition of the Ras-MAPK pathway results in AXIN1 loss in CRC.

In this study, we deciphered cellular mechanisms underlying AXIN1 loss by MEK1/2 inhibition in CRC. We observed that both MEK1/2 and GSK3B inhibitors strongly reduce AXIN1 levels across multiple CRC models. Inhibition of MEK1/2, but not GSK3B, causes a reduction of *AXIN1* transcript levels. While GSK3B inhibition results in rapid protein degradation of AXIN1, targeting of MEK1/2 alters neither protein stability nor post-translational modifications of AXIN1. Instead, MEK1/2 inhibition represses global translation via an mTOR dependent mechanism, which is sufficient to cause AXIN1 loss. Concordantly, mTOR inhibitors phenocopy the effect of MEK1/2 inhibitors on AXIN1 levels. Our study demonstrates that AXIN1 protein homeostasis is critically controlled by Ras-MAPK signaling at the level of protein synthesis, and that its perturbation by MEK1/2 inhibitors result in AXIN1 loss.

## RESULTS

### AXIN1 loss after MEK1/2 and GSK3B inhibition shows distinct characteristics

We previously observed that pharmacological inhibition of MEK1/2 and GSK3B caused a reduction of AXIN1 protein levels in CRC cell lines (*7*). To determine if this effect is observed across a broader range of CRC models with diverse genetic backgrounds, we treated eight CRC cell lines and two patient-derived CRC organoid lines with the MEK1/2 inhibitor trametinib or the GSK3B inhibitor CHIR90221 (Fig. 1A). In all CRC cell and organoid lines, trametinib efficiently inhibits Ras signaling, as evidenced by reduced phospho-ERK1/2 levels. AXIN1 protein levels were decreased upon MEK1/2 inhibition in all CRC lines with the exception of SNUC2A, and the effect size varied between individual lines. Similarly, GSK3B inhibition caused AXIN1 loss in most cell lines, and the reduction was more pronounced compared to MEK1/2 inhibition (Fig. 1A). Likewise, both MEK1/2 and GSK3B inhibitors reduced AXIN1 levels in patient-derived CRC organoids (Fig. 1A). These results demonstrate that loss of AXIN1 is a general consequence of MEK1/2 and GSK3B inhibition in different CRC models.

**Fig. 1:**
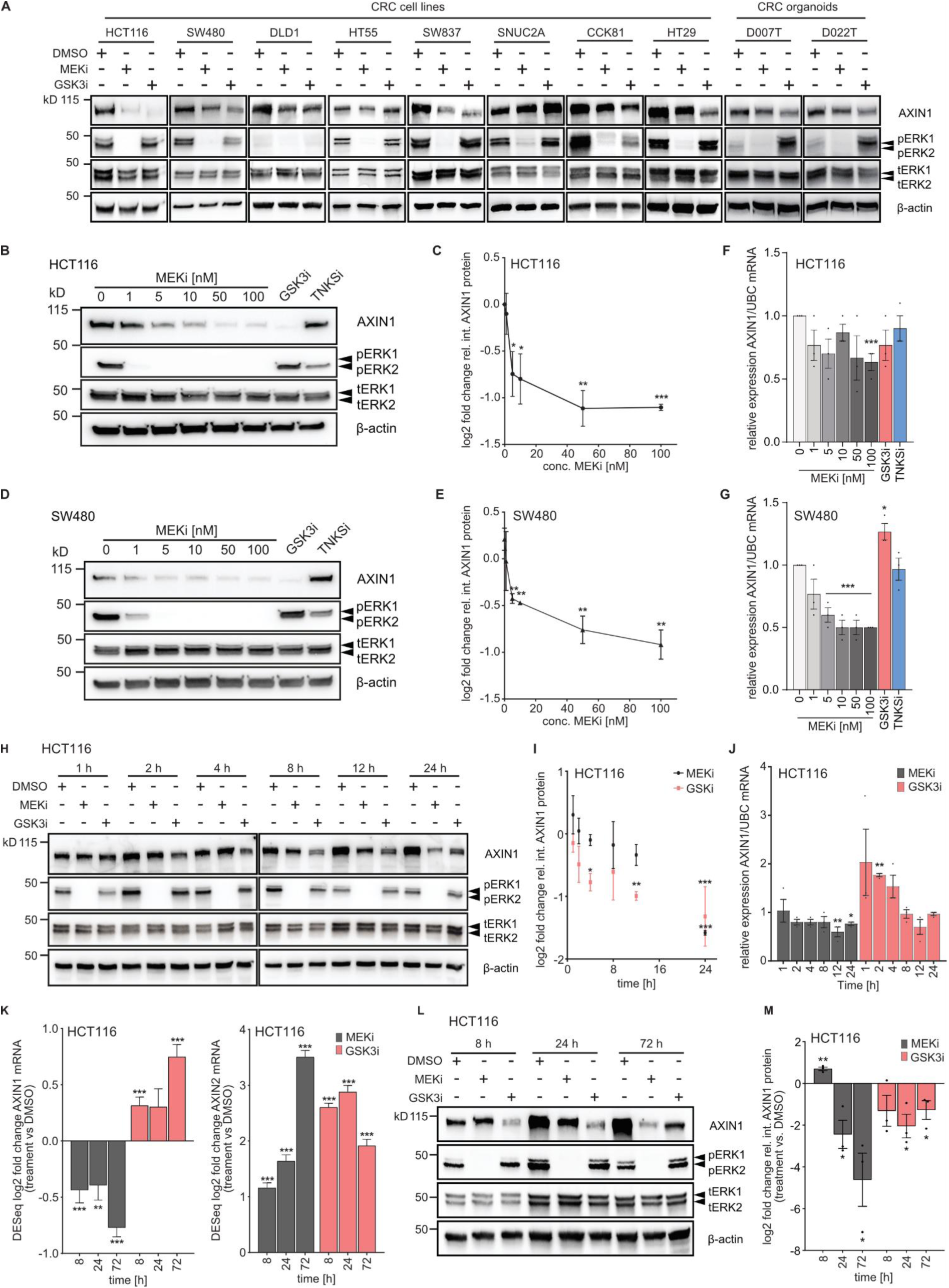
MEK1/2 and GSK3B inhibitors induce AXIN1 loss in different models of colorectal cancer. **A**, AXIN1 protein levels after MEK1/2 and GSK3B inhibition in a panel of eight CRC cell lines and two patient-derived organoid lines. Drug treatment was performed for 24 h for CRC cell lines and 72 h for organoids. MEKi: 100 nM trametinib, GSKi: 10 μM CHIR-99021 **B-C,** Concentration-dependent effect of MEK1/2 inhibition on AXIN1 protein levels in HCT116 after treatment for 24 h. TNKSi: 10 μM XAV979. Representative immunoblot image is shown in (B) and quantification of replicates is shown in (C). **D-E,** Concentration-dependent effect of MEK1/2 inhibition on AXIN1 protein levels in SW480 after treatment for 24 h. Representative immunoblot image is shown in (D) and quantification of replicates is shown in (E). **F,** Concentration-dependent effect of MEK1/2 inhibition on *AXIN1* mRNA levels in HCT116 after treatment for 24 h. **G,** Concentration-dependent effect of MEK1/2 inhibition on *AXIN1* mRNA levels in SW480 after treatment for 24 h. **H-I,** Time-dependent effect of MEK1/2 and GSK3B inhibition on AXIN1 protein levels in HCT116. Representative immunoblot image is shown in (H) and quantification of replicates is shown in (I). **J,** Time-dependent effect of MEK1/2 inhibition on *AXIN1* mRNA levels in HCT116. **K,** Time-dependent effect of MEK1/2 and GSK3B inhibition on *AXIN1* and *AXIN2* transcript levels in HCT116, analyzed by RNAseq. Cells were treated for the indicated time periods with the inhibitors. Statistical analysis was performed using DESeq2. Data from three independent experiments are presented as log2 fold change ± lfcSE as determined by DESeq2. **L-M,** Time-dependent effect of MEK1/2 and GSK3B inhibition on AXIN1 protein levels. Representative immunoblot image is shown in (L) and quantification of replicates is shown in (M). If not otherwise stated, data from three independent experiments are presented as mean ± SEM \**p* < 0.05, \*\**p* < 0.01, \*\*\**p* < 0.001, two-tailed Student’s t-test. Drug concentrations were as follows: MEKi - 100 nM trametinib, GSK3Bi - 10 μM CHIR-99021, TNKSi - 10 μM XAV979.

Next, we characterized the concentration-dependent effect of MEK1/2 inhibition on AXIN1 levels in the two CRC cell lines HCT116 and SW480 (Fig. 1B-E). We observed a significant reduction of AXIN1 protein levels starting from 5 nM of trametinib in both cell lines. Concurrent measurement of *AXIN1* transcript levels demonstrated a reduction down to 63% in HCT116 and 50% in SW480 at 100 nM trametinib, while GSK3B inhibition did not significantly reduce *AXIN1* mRNA levels (Fig. 1F-G). Transcriptional induction of *AXIN2*, indicative of Wnt activation, was already observed at low concentrations of the MEK1/2 inhibitor (Fig. S1A-B). We then assessed the temporal dynamics of AXIN1 loss upon MEK1/2 and GSK3B inhibition in HCT116 cells. Compared to GSK3B inhibition, which reduced AXIN1 protein levels significantly between 8-12 h, the effect of MEK1/2 inhibitors was more pronounced at later time points between 12-24 h, indicating different modes-of-action (Fig. 1H-I). Significant reduction of phospho-ERK1/2 levels was observed already 1 h after MEK1/2 inhibition, suggesting that AXIN1 loss occurs after a latency period upon inhibition of Ras-MAPK signaling (Fig. 1H). Concurrent measurement of *AXIN1* transcript levels showed a mild reduction after 12 and 24 h after MEK1/2 inhibition, whereas transcript levels initially increase after GSK3B inhibition and then gradually return to their initial values (Fig. 1J). Since AXIN1 is localized in the nucleus and cytoplasm, we sought to determine if AXIN1 is preferentially reduced in a specific subcellular compartment. To this end, we performed subcellular fractionation after MEK1/2 and GSK3B inhibition in HCT116 cells, separating the nuclear and cytoplasmic fraction. Results of this experiment demonstrate that AXIN1 protein is mainly detected in the cytoplasm, but its loss upon MEK1/2 inhibition occurs in both fractions (Fig. S1C). Next, we asked if transcriptomic changes induced by MEK1/2 and GSK3B inhibition are fundamentally different and persist beyond 24 h of treatment. We performed RNAseq of HCT116 and SW480 cells treated for 8, 24 and 72 h with MEK1/2 and GSK3B inhibitors, and determined protein levels of AXIN1 in parallel. RNAseq results revealed distinct transcriptomic shifts over time and strong differences between MEK1/2 and GSK3B inhibition in both cell lines, as shown by principal component analysis (Fig. S1D-E). In line with our previous results, *AXIN1* transcript levels were progressively reduced upon MEK1/2 inhibition, whereas they were increased upon GSK3B inhibition (Fig. 1K). *AXIN2* transcript levels increased over time after MEK1/2 inhibition, whereas their induction peaked at 24 h after GSK3B inhibition. A similar change of *AXIN1* and *AXIN2* expression was observed in SW480 cells (Fig. S1F-G), with the exception that GSK3B inhibition failed to induce *AXIN2* expression. Parallel measurement of AXIN1 protein levels in HCT116 cells showed an enhanced repression over time after MEK1/2 inhibition, while AXIN1 levels did not further decrease between 24 h and 72 h of GSK3B inhibition (Fig. 1L-M). Comparison of transcript and protein levels after MEK1/2 inhibition showed that AXIN1 loss was stronger at the protein than at the transcript level (mRNA reduction to 58% mRNA level of DMSO control versus protein reduction to 4% protein level of DMSO control at 72 h; Fig. 1K, M). In summary, these results show that MEK1/2 and GSK3B inhibition both cause repression of AXIN1 protein levels, but with distinct kinetics. Furthermore, MEK1/2, but not GSK3B inhibition, represses transcription of AXIN1 mRNA. Nevertheless, reduced AXIN1 transcript levels do not fully account for the strong repression at the protein level after MEK1/2 inhibition.

### AXIN1 loss after MEK inhibition is not mediated by active protein degradation

Cellular levels of AXIN1 are extensively regulated by post-translational modifications and subsequent changes in protein stability, with several E3 ubiquitin ligases and deubiquitinases reported to be involved in this process. To understand if MEK1/2 inhibition induces protein degradation of AXIN1, we first performed cycloheximide (CHX) chase assays which allowed us to monitor AXIN1 stability after blockage of de novo protein synthesis and to calculate the protein half-life. AXIN1 protein levels gradually decrease over time, with a half-life of 14.4 h in HCT116 cells (Fig 2A). This finding is in line with data from a proteome-wide mapping of protein stabilities, which determined a half-life of 13 h for AXIN1 in the same cell line (*24*). Next, we assessed the stability of AXIN1 upon additional pharmacological inhibition of tankyrases, GSK3B and MEK1/2 (Fig. 2B-D). Cells were pretreated with the inhibitors and CHX was added for a total of 12 h. We selected different pretreatment periods for the individual inhibitors to ensure that no changes in AXIN1 are induced prior to addition of CHX. In line with the important role of tankyrases in PARsylation-mediated degradation of AXIN1, treatment with the tankyrase inhibitor XAV979 stabilized AXIN1 in the cycloheximide chase assay (Fig. 2B). In contrast, AXIN1 degradation was strongly induced upon GSK3B inhibition and CHX treatment (Fig. 2C). Upon MEK inhibitor treatment, however, no significant change in the kinetics of AXIN1 loss was observed upon blockage of de novo translation (Fig. 2D).

**Fig. 2:**
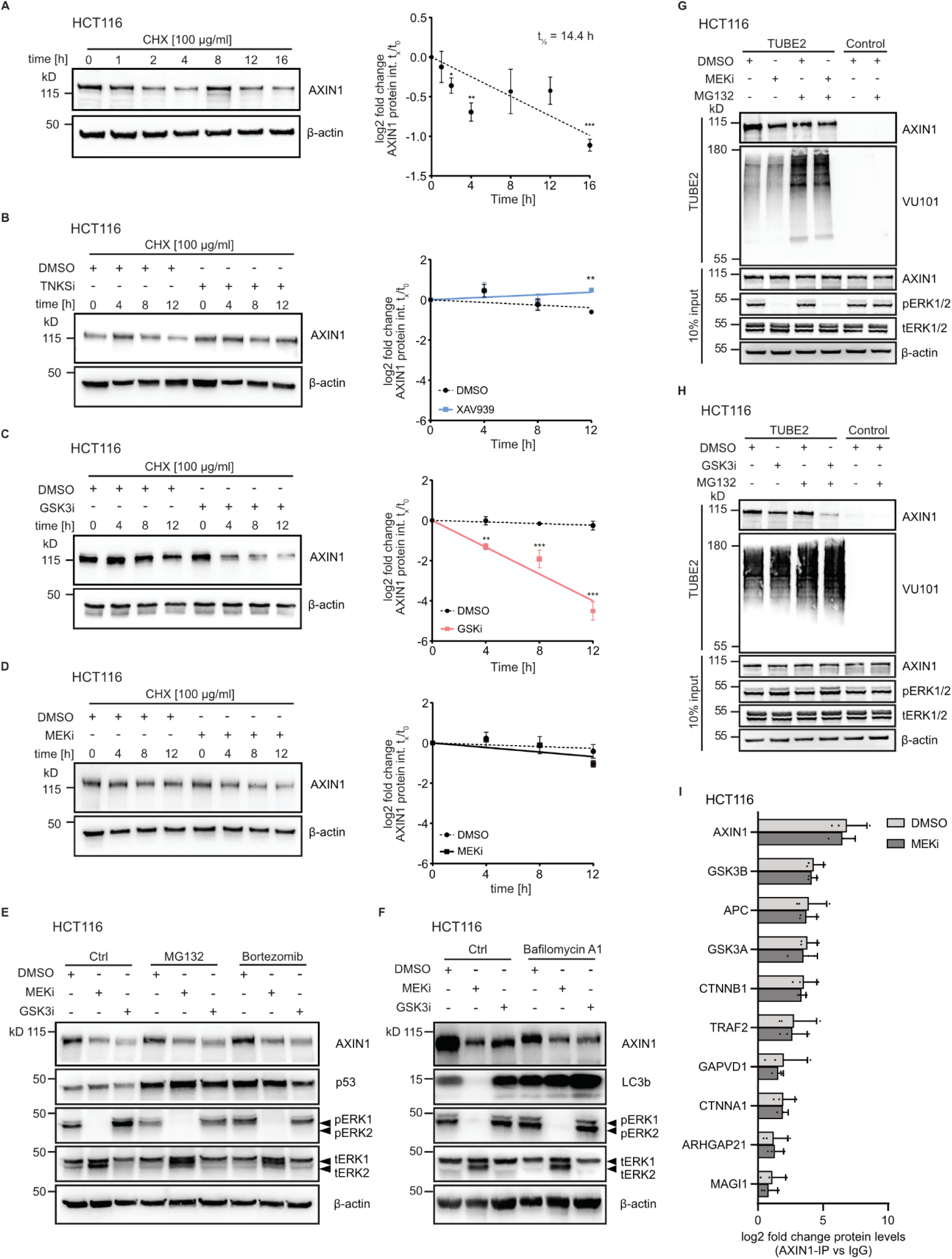
Loss of AXIN1 after MEK1/2 inhibition is not mediated by changes in protein stability. **A**, Measurement of AXIN1 protein half-life in HCT116 by cycloheximide chase assay. Cells were treated with cycloheximide (CHX) and protein levels of AXIN1 were determined at indicated time points by immunoblot. Representative immunoblot image (left) and quantification of replicates (right) are shown. **B,** Measurement of AXIN1 half-life under treatment with a tankyrase inhibitor by CHX chase assay. Cells were treated with 10 μM XAV979 (TNKSi) or DMSO for 8 h, followed by co-treatment with CHX for the indicated time periods. Representative immunoblot image (left) and quantification of replicates (right) are shown. **C,** Measurement of AXIN1 half-life under treatment with a GSK3B inhibitor by CHX chase assay. Cells were pretreated for 1 h with 10 μM CHIR-99021 (GSK3i) or DMSO, followed by co-treatment with CHX for the indicated time periods. Representative immunoblot image (left) and quantification of replicates (right) are shown. **D,** Measurement of AXIN1 half-life under treatment with a MEK1/2 inhibitor by CHX chase assay. Cells were pretreated for 8 h with 100 nM trametinib (MEKi) or DMSO, followed by co-treatment with CHX for the indicated time periods. Representative immunoblot image (left) and quantification of replicates (right) are shown. A-D: Data from three independent experiments are presented as mean ± SEM *p < 0.05, **p < 0.01, ***p < 0.001, two-tailed Student’s t-test. **E,** Proteasomal inhibition does not prevent MEK1/2 and GSK3B inhibitor-induced AXIN1 loss. Cells were treated with MEKi and GSK3i for 16 h, followed by 4 h of co-incubation with 10 μM MG132 or 10 μM bortezomib. Accumulation of TP53 expression indicates cellular stress induced by proteasome inhibition. **F,** Lysosomal inhibition does not prevent MEK1/2 and GSK3B inhibitor-induced AXIN1 loss. Cells were treated with MEKi and GSK3i for 8 h, followed by 24 h of co-incubation with 100 nM bafilomycin A1. Accumulation of LC3b indicates efficient inhibition of lysosomal function. **G-H,** Ubiquitin-affinity precipitation does not show increased polyubiquitination of AXIN1 upon MEKi or GSK3i. Cells were treated for 4 h with 100 nM trametinib (MEKi, G) or 30 min with 10 μM CHIR-99021 (GSKi, H), or co-treated with 20 μM MG132, followed by precipitation with pan-selective tandem ubiquitin binding entities (TUBEs). Ubiquitination is detected by the pan-ubiquitin recognizing antibody VU101. Representative images from three replicates are shown. Drug concentrations were as follows: MEKi: 100 nM trametinib, GSK3Bi: 10 μM CHIR-99021, TNKSi: 10 μM XAV979. **I**, Protein-protein interactions of AXIN1 with destruction complex partners do not change after MEKi. Cells were treated for 4 h with 100 nM trametinib, followed by Co-IP with anti-AXIN1 antibody or IgG control, and mass spectrometry analysis. Ten most enriched proteins after IP with anti-AXIN1 antibody are shown. Data from three independent experiments are presented as mean ± SD.

Next, we assessed if inhibition of proteasomal or lysosomal degradation pathways would interfere with AXIN1 turnover after MEK1/2 or GSK3B inhibition. Pre-treatment with MEK1/2 or GSK3B inhibitors, followed by inhibition of the proteasome using either MG132 or bortezomib did not prevent loss of AXIN1 loss despite accumulation of p53, which is indicative of compromised proteasome function (*25*) (Fig. 2E). Since AXIN1 turnover can be controlled by the autophagy-lysosome pathway (*26*), we tested if inhibition of lysosomal degradation by bafilomycin A1 affects MEK1/2 or GSK3B-induced AXIN1 loss. Bafilomycin A1 led to elevated LC3b levels indicative of perturbed lysosomal function, and surprisingly caused a reduction of AXIN1 protein levels. However, inhibition of lysosomal degradation did not prevent loss of AXIN1 upon MEK1/2 or GSK3B inhibition (Fig. 2F). Proteins undergoing proteasomal degradation are marked by polyubiquitination. To determine if MEK1/2 inhibition changes AXIN1 ubiquitination, we performed ubiquitin-affinity precipitation using pan-selective tandem ubiquitin binding entities (TUBEs). To detect changes in polyubiquitination prior to subsequent protein degradation, we selected short treatment periods during which cellular AXIN1 levels were not yet changed by the inhibitors (4 h for trametinib and 30 min for CHIR99021). Results of these experiments show that global protein ubiquitination is increased upon proteasome inhibition by MG132, as determined by an antibody that recognizes a broad spectrum of ubiquitin linkages (VU101) (Fig. 2G-H). AXIN1 polyubiquitination was mildly reduced upon MEK1/2 inhibition under native conditions, but not when cells were treated with MG132 (Fig. 2G). In contrast, GSK3B led to a pronounced reduction of ubiquitinated AXIN1, particularly after co-treatment with MG132 (Fig. 2H). While the basis for this effect remains unclear, we concluded that AXIN1 is not targeted for degradation by the ubiquitin-proteasome system.

To determine if MEK1/2 and GSK3B inhibition induce changes in protein-protein interactions of AXIN1 and GSK3B, we performed co-immunoprecipitation (Co-IP) with antibodies targeting both proteins, followed by mass spectrometry analysis. Treatment conditions were the same as for the ubiquitin-affinity precipitation experiments, as we aimed to identify changes in interaction that occur prior to AXIN1 loss. A clear enrichment of AXIN1 and other major interaction partners within the destruction complex was seen after AXIN1 and GSK3B Co-IP (Fig. 2I, Fig. S2A). However, we did not observe any significant changes in protein-protein interactions of destruction complex members upon MEK1/2 inhibition (Fig. 2I, S2A-C). This result was confirmed by immunoblot analysis following AXIN1 and GSK3B Co-IP, showing no change of interaction between GSK3B, AXIN1 and beta-catenin during the selected incubation periods (Fig. S2D-E). After GSK3B inhibition, we observed an overall increase of protein-protein interactions for AXIN1, and a stronger association of AXIN1 with beta-catenin (Fig. S2F-I). Analysis of different post-translational modifications of AXIN1 only identified phosphorylation with high confidence. However, AXIN1 phosphorylation was not changed upon MEK1/2 inhibition (Fig. S2J). In summary, using different methodological approaches, we show that MEK1/2 and GSK3B inhibition induce AXIN1 loss by distinct mechanisms. While GSK3B inhibitors cause AXIN1 degradation, MEK1/2 inhibitors do not change protein stability, interactions with destruction complex members or phosphorylation of AXIN1.

### Tankyrase inhibition partially prevents AXIN1 loss upon MEK1/2 and GSK3B inhibition

PARsylation by tankyrases is an essential post-translational modification that targets AXIN1 for subsequent ubiquitination and degradation by the E3 ligase RNF146, thereby regulating protein homeostasis of AXIN1 (*17*, *27*). We hypothesized that pharmacological inhibition of tankyrase-mediated AXIN1 degradation may modify the effect of MEK1/2 and GSK3B inhibitors on AXIN1. To this end, we pretreated HCT116 cells with the two inhibitors for 8 h, followed by co-treatment with XAV979 for 24 h. Under control conditions with DMSO, tankyrase inhibition led to an increase of AXIN1 levels (Fig. 3A). AXIN1 levels were strongly reduced upon MEK1/2 and GSK3B inhibition, and co-treatment with XAV979 partially prevented this effect. However, prevention of AXIN1 loss by XAV979 was significantly stronger for GSK3B inhibitor versus MEK1/2 inhibitor treatment. Based on this observation, we sought to determine if MEK or GSK3B inhibition alters AXIN1 levels by directly changing the expression of tankyrases. We measured protein levels of TNKS1 and TNKS2 at 8, 12 and 24 h after addition of GSK3B and MEK1/2 inhibitors (Fig. 3B). Our results show that after 24 h, GSK3B inhibition induced protein levels of both tankyrases, whereas MEK1/2 inhibition clearly reduced them. Parallel measurement of *TNKS1* and *TNKS2* transcript levels showed no effect of MEK1/2 inhibition, whereas *TNKS1* mRNA was reduced at all time points after GSK3B inhibition (Fig. 3C-D).

**Fig. 3:**
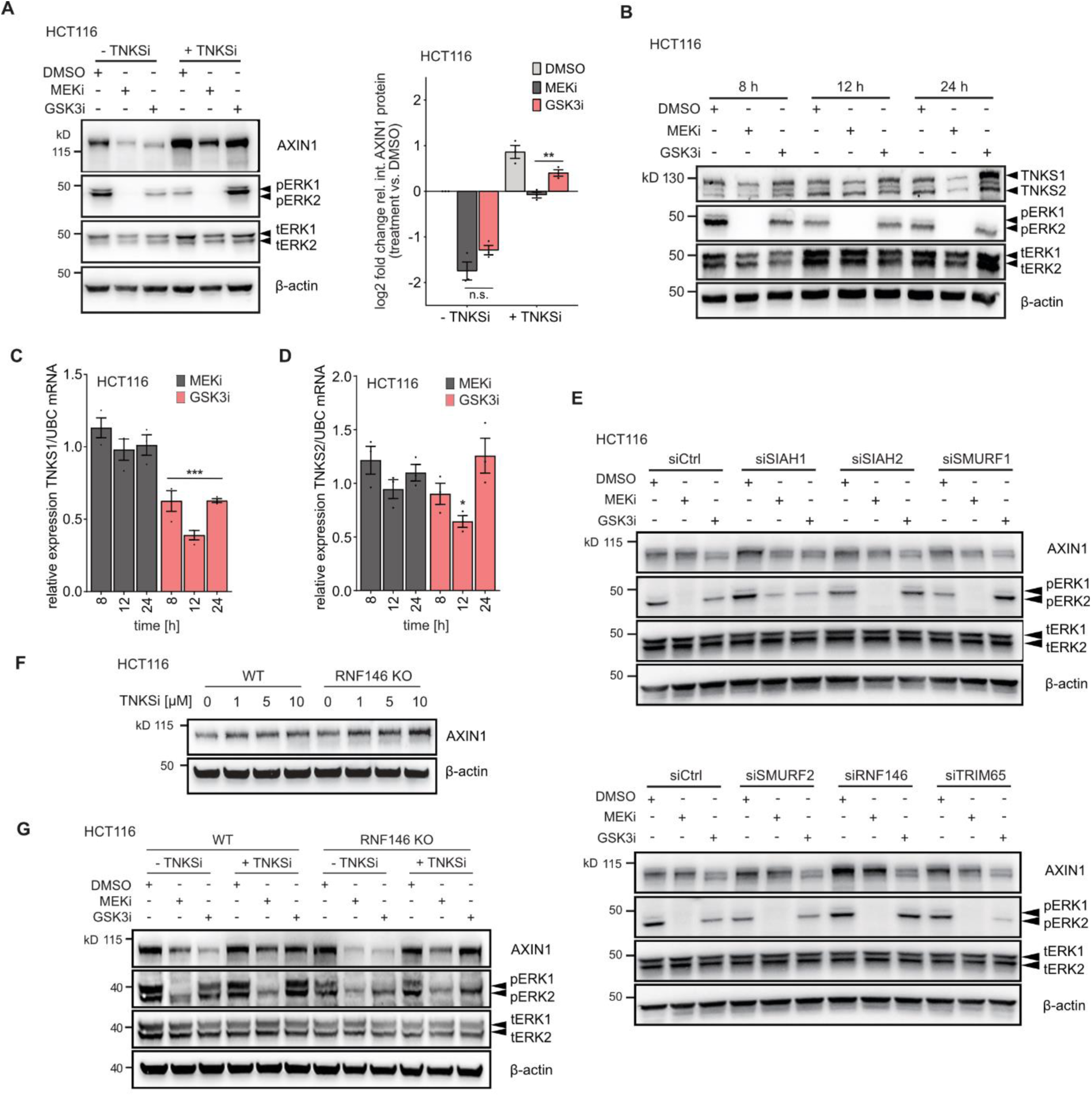
Tankyrase inhibition partially prevents AXIN1 loss upon MEK1/2 and GSK3B inhibition. **A**, Effect of tankyrase inhibition on AXIN1 loss upon MEK1/2 and GSK3B inhibition. HCT116 cells were pretreated with 100 nM trametinib (MEKi), 10 μM CHIR-99021 (GSK3i) or DMSO for 8 h, followed by co-treatment with 10 μM XAV979 (TNKSi) for 24 h. Representative immunoblot image (left) and quantification of replicates (right) are shown. **B,** Effect of MEK1/2 and GSK3B inhibition on TNKS1 and TNKS2 protein levels after indicated treatment periods. A representative image of three replicates is shown. **C-D,** Effect of MEK1/2 and GSK3B inhibition on TNKS1 and TNKS2 transcript levels after indicated treatment periods. Expression of TNKS1/2 is compared to the DMSO control of the same time points. Data from three independent experiments are presented as mean ± SEM \**p* < 0.05, \*\*\**p* < 0.001, two-tailed Student’s t-test. **E,** Effect of RNAi-mediated knockdown of selected E3 ubiquitin ligases on AXIN1 loss upon MEK1/2 and GSK3B inhibition. Cells were treated with siRNA pools for 48 h, and then incubated with the inhibitors for 24 h. A representative image of three replicates is shown. **F**, CRISPR-mediated RNF146 knockout enhances TNKSi-mediated AXIN1 stabilization in HCT116 cells. Isogenic cell lines were treated for 24 h with the indicated drug concentrations. **G**, RNF146 knockout in combination with TNSKi does not prevent AXIN1 loss upon MEK1/2 inhibition. HCT116 cell lines were co-treated with 5 μM XAV979 plus trametinib (MEKi), CHIR-99021 (GSK3i) or DMSO for 24 h. A-B, E-G: Representative images from three independent replicates are shown. Drug concentrations were as follows: MEKi - 100 nM trametinib, GSK3Bi - 10 μM CHIR-99021.

Since tankyrase inhibition increased basal levels of AXIN1 and partially prevented MEK1/2 and GSK3B inhibitor-induced loss, we sought to determine if functional depletion of RNF146 and other AXIN1 targeting E3 ubiquitin ligases would phenocopy this effect. We selected the E3 ubiquitin ligases SIAH1, SIAH2, RNF146, SMURF1, SMURF2 and TRIM65 for knockdown using siRNAs. RNAi successfully depleted expression in all cases as determined by qPCR (Fig. S3A). Basal levels of AXIN1 were increased upon knockdown of specific E3 ubiquitin ligases such as RNF146, underlining their role in controlling AXIN1 protein homeostasis (Fig. 3E). Concurrent measurement of transcript levels demonstrates that only SMURF2 knock-down mildly increased *AXIN1* mRNA expression (Fig. S3B). However, knockdown of none of the E3 ubiquitin ligases prevented AXIN1 loss after 24 h treatment with MEK1/2 or GSK3B inhibitors (Fig. 3E).

To confirm these results using a complementary approach, we performed CRISPR-mediated knockout of RNF146 in HCT116 using a previously validated sgRNA sequence (*28*). RNF146 was selected based on its strong effect on basal AXIN1 levels and the functional cooperation with TNKS1/2 in controlling AXIN1 turnover. We observed slightly increased basal AXIN1 levels in RNF146 knockout cells (Fig. 3F). This knockout cell line was also more responsive to tankyrase inhibitor-mediated AXIN1 stabilization, as AXIN1 levels were higher across different XAV979 concentrations (Fig. 3F). We then tested the effect of GSK3B and MEK1/2 inhibitors on AXIN1 levels in these two cell lines, in the presence and absence of XAV979 (Fig. 3G). Similar to siRNA-mediated knockdown of RNF146 (Fig 3E), knockout of RNF146 did not prevent GSKB and MEK1/2 inhibitor-induced AXIN1 loss. In line with our previous results, co-treatment of the two inhibitors with XAV979 partially prevented AXIN1 loss. This effect was again stronger in cells treated with GSK3B than MEK1/2 inhibitors, but not different between wild-type and RNF146 knockout cells. Together, these results suggest that inhibition of AXIN1 PARsylation attenuates the loss of AXIN1 upon GSK3B and MEK1/2 inhibition, with the effect being significantly weaker for the latter. While depletion of specific E3 ubiquitin ligases increases basal AXIN1 levels, it does not prevent MEK1/2 inhibitor-induced AXIN1 loss. These results indicate that MEK1/2 inhibitors induce AXIN1 loss by mechanisms that can partially override increased protein stability.

### Effect of APC truncation on MEK1/2 inhibitor-induced Wnt activation and AXIN1 loss

We previously showed that APC truncations increase MEK-inhibitor induced Wnt activation in isogenic CRC cell lines (*7*). We hypothesized that this effect might be caused by altered dynamics of AXIN1 loss. To address this question, we used two isogenic CRC models harboring APC mutations or deletions (Fig. 4A). First, we used an isogenic HCT116 cell line that harbors two different APC truncations in a mutational hot-spot locus. This cell line was generated using CRISPR/Cas9, and truncation of APC was previously confirmed by immunoblot and sequencing (*29*). In cell lines harboring the APC truncations, we observed an increased basal and MEK1/2 inhibitor-induced activation in Wnt signaling, as shown by elevated *AXIN2* transcript levels (Fig. 4B). Interestingly, GSK3B inhibitor-mediated Wnt activation was abolished upon APC truncation, supporting the notion that the both inhibitors act through distinct mechanisms to stimulate Wnt signaling. We then assessed basal protein levels of AXIN1 and observed a strong decrease in APC truncated versus wild-type cell lines (Fig. 4C). Nevertheless, treatment with both the MEK1/2 and the GSK3B inhibitor led to a further reduction of AXIN1 protein levels in APC truncated cells (Fig. 4D).

**Fig. 4:**
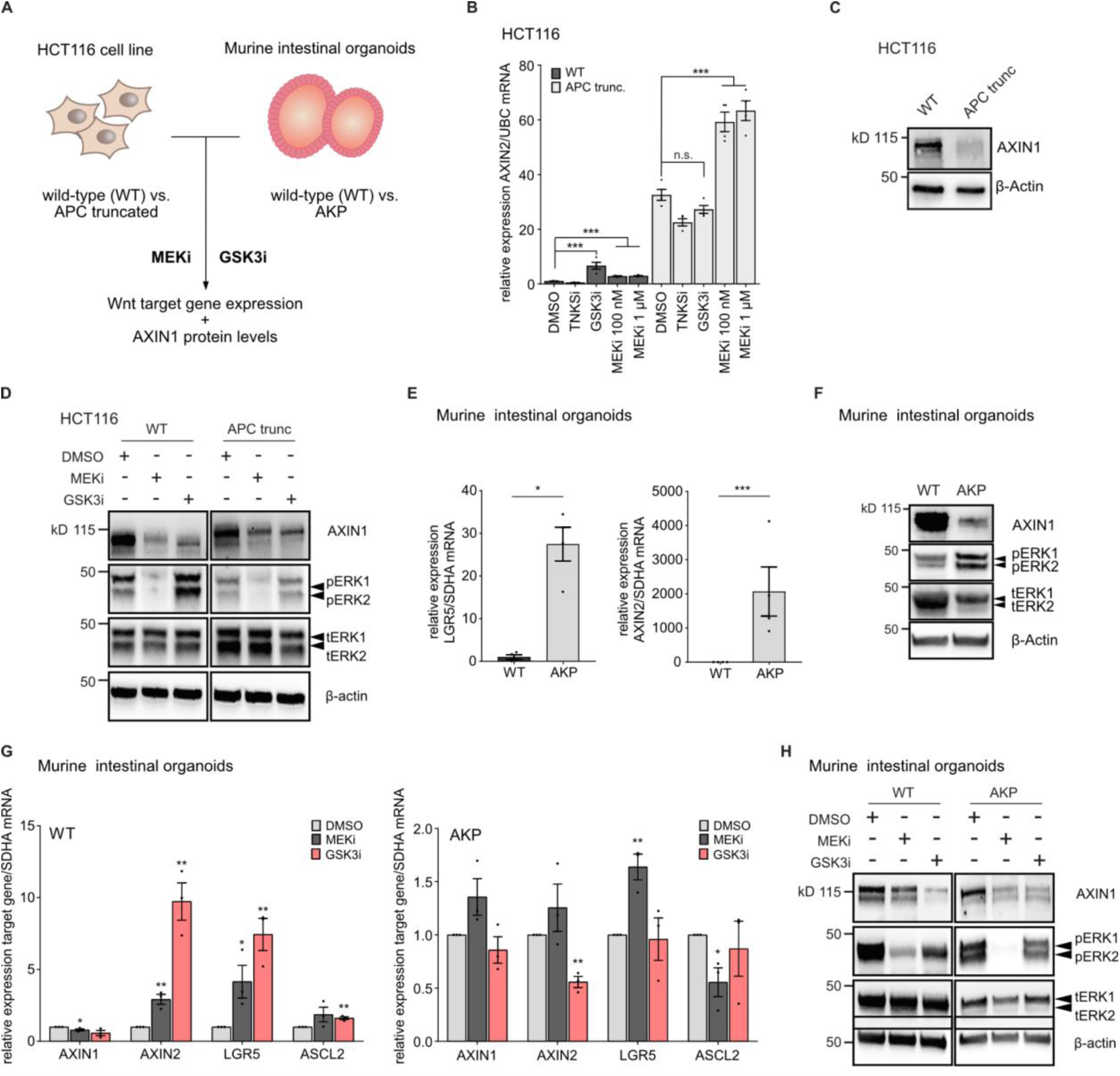
APC truncations modulate AXIN1 levels andMEK1/2 and GSK3B inhibitor-induced Wnt activation. **A**, Schematic image of CRC models and treatment conditions. Isogenic HCT116 cell lines with APC truncation and isogenic intestinal organoids from Apc^fl/fl^, Kras^G12D/+^, Trp53^fl/fl^ C57B6 mice are treated with 100 nM trametinib (MEKi) or 10 μM CHIR-99021 (GSKi), followed by profiling of Wnt target gene expression and AXIN1 levels. **B**, APC truncations in HCT116 enhance basal and MEK1/2 inhibitor-induced Wnt activation. **C**, Basal protein level of AXIN1 is reduced in APC truncated HCT116 cells. **D,** Effect of MEK1/2 and GSK3B inhibitor on AXIN1 protein level in wild-type (WT, left) and APC truncated (right) HCT116. Representative immunoblots from three independent replicates are shown in C and D. **E,** Expression of Wnt target genes are increased in AKP murine intestinal organoids (AKP) compared to wild-type intestinal organoids (WT). **F**, Basal protein levels of AXIN1 are reduced and (phospho-)ERK1/2 levels are increased in AKP versus WT organoids. **G**, Differential induction of Wnt target gene expression in WT (left) and AKP intestinal organoids (right) by MEKi and GSK3i after 72 h treatment. **H**, MEK1/2 and GSK3B inhibitors reduce AXIN1 protein levels in wild-type and AKP murine intestinal organoids after 72 h. B, E, G: Data from at least three independent experiments are presented as mean ± SEM *p < 0.05, **p < 0.01, ***p < 0.001, two-tailed Student’s t-test. Drug concentrations were as follows: MEKi - 100 nM trametinib, GSK3i - 10 μM CHIR-99021.

To validate these findings in an independent model system, we used mouse intestinal organoids from wild-type C57B6 mice and C57B6 with Apc^fl/fl^, Kras^G12D/+^ and Trp53^fl/fl^ background (AKP) (*30*). Compared to the wild-type counterparts, transcript levels of the Wnt target genes and intestinal stem cell markers AXIN2 and LGR5 were strongly induced in AKP organoids (Fig. 4E). Similar to our observation in isogenic cell lines, loss of Apc markedly reduced basal AXIN1 protein levels, while the Kras^G12D/+^ mutation enhanced Ras-MAPK pathway activity, as evidenced by elevated phospho-ERK1/2 levels (Fig. 4F). Both MEK1/2 and GSK3B inhibition stimulated mRNA expression of *AXIN2* and *LGR5* in wild-type organoids, and GSK3B inhibition also led to an increase of *ASCL2* expression. However, in AKP organoids, only MEK1/2 inhibition increased *LGR5* expression significantly, whereas the GSK3B inhibitor did not induce expression of Wnt target genes (Fig. 4G). Despite the strongly reduced basal AXIN1 levels in AKP organoids, both MEK1/2 and GSK3B inhibition led to a further decrease in AXIN1 protein levels (Fig. 4H). In summary, both isogenic model systems show that APC truncations or deletions result in strongly reduce basal AXIN1 levels and an increase in Wnt target gene expression. However, AXIN1 loss upon MEK1/2 or GSK3B inhibition was not affected by APC truncations or deletions.

### MEK1/2 inhibition represses global protein synthesis via mTOR

Our previous experiments indicate that loss of AXIN1 by MEK1/2 inhibition, as opposed to GSK3B inhibition, is not mediated by protein destabilization. Furthermore, transcriptional repression of AXIN1 by MEK1/2 inhibition does not fully explain the strong reduction of AXIN1 observed at the protein level (Fig. 1K-M). Therefore, we assumed that additional mechanisms exist by which MEK1/2 inhibition reduces AXIN1 levels. Since Ras-MAPK signaling critically controls protein synthesis, we hypothesized that MEK1/2 inhibition may reduce translation of *AXIN1* mRNA. To address this question, we first performed polysome profiling experiments to measure the rate of global protein synthesis after MEK1/2 and GSK3B inhibition in HCT116 cells. Our results show that MEK1/2 inhibition for 24 h resulted in a 46% reduction of the polysomal area under the curve, a proxy for the proportion of ribosomes that are actively engaged in translation (Fig. 5A). In contrast, GSK3B inhibition slightly increased global translation compared to the DMSO control, although this effect was not statistically significant.

**Fig. 5:**
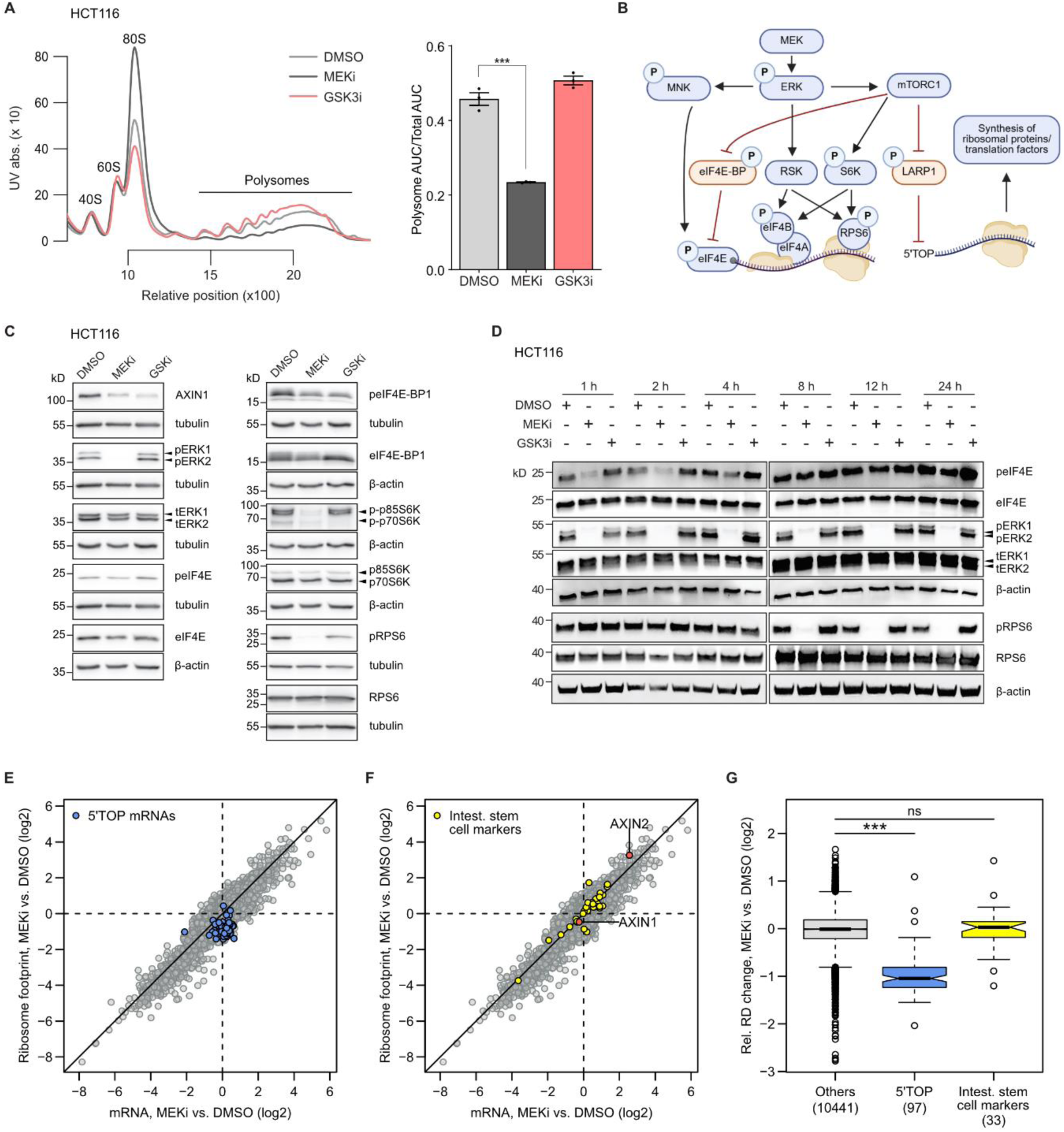
MEK1/2 inhibition represses global and mRNA-specific protein biosynthesis. **A**, Polysome profiling was used to measure global protein biosynthesis in HCT116 after treatment with 100 nM trametinib (MEKi) or 10 μM CHIR-99021(GSKi) for 24 h. For quantification, the area under the curve (AUC) of the polysomal part was divided by the total area. Data from three independent experiments are presented as mean ± SD, and differences were tested using a two-tailed Student’s t-test (***p < 0.001). **B,** Scheme of signalling events downstream of MEK activation that affect translation initiation. Positive regulators of translation are depicted in blue, negative regulators in orange. **C,** Effect of MEKi and GSKi (as in panel A) on the phosphorylation of effector proteins downstream of MEK as detected by Western blotting. **D,** Time-resolved analysis of eIF4E and RPS6 phosphorylation upon MEK1/2 and GSK3B inhibition. A representative image of three independent replicates is shown. **E,** RiboSeq was performed in HCT116 cells after treatment with 100 nM trametinib (MEKi) for 24 h. 5’TOP mRNAs are highlighted in blue. **F,** RiboSeq results as in panel E with Wnt-associated intestinal stemness markers (yellow) and AXIN1/2 (orange). **G,** Relative changes in ribosome densities (RD) were calculated from MEKi-induced changes in ribosome footprint levels divided by changes in input levels and shown as box-and-whisker plot. Differences were tested using a two-sided Wilcoxon rank sum test (***p < 0.001).

We then explored if global translation repression by MEK1/2 inhibition might also affect the protein abundance of other Wnt pathway components. To this end, we made use of our previously published global proteomic profiling datasets of HCT116 and SW480 cells treated with 100 nM trametinib for 24 h (*31*). Due to their low abundance, AXIN1 and many other Wnt pathway components were not detectable in the datasets (Fig. S4A). However, for components that were detectable, no significant changes in protein abundances were observed after MEK1/2 inhibition. These results indicate that repression of protein synthesis by MEK1/2 inhibition has a particularly strong effect on AXIN1 levels.

The Ras-MAPK pathway controls translation via three main routes: the MNK and RSK kinases as well as mTOR (see overview Fig. 5B) (*32*). To mechanistically understand which of these pathways mediates reduced protein synthesis upon MEK1/2 inhibition, we profiled the phosphorylation status of eIF4E, eIF4E-BP1, S6K1 and RPS6. After 24 h, phosphorylation of eIF4E-BP1 and the p70 isoform of S6K1 was reduced by both MEK1/2 and GSK3B inhibition (Fig. 5C). This is in line with reports that GSK3B can also phosphorylate eIF4E-BP1 (*33*) and p70S6K (*34*). Phosphorylation of the p85 isoform of S6K1 was only affected by MEK1/2 inhibition, and RPS6 phosphorylation was decreased more strongly by MEK1/2 inhibition than by GSK3B inhibition. Kinetic analysis demonstrates that eIF4E phosphorylation was only transiently reduced by MEK1/2 inhibition within the first 1-4 h, after which it returned to basal levels. In contrast, phosphorylation of RPS6 was constantly reduced after 8-12 h (Fig. 5D). These results indicate that neither eIF4E phosphorylation nor inactivation by eIF4E-BP1 contribute to the translational suppression observed after 24 h of MEK1/2 inhibition. However, the pronounced loss of p85 S6K1 phosphorylation is specific for MEK1/2 inhibition and may thus play a role in dampening protein synthesis. Phosphorylation of LARP1 could not be assessed due to the lack of an adequate antibody. Unless it is phosphorylated by mTORC1, LARP1 represses translation of a subset of mRNAs that bear a 5’TOP motif (*35*). These mRNAs encode for ribosomal proteins as well as several translation initiation and elongation factors (*36*). Therefore, specific regulation of 5’TOP mRNA translation by mTOR represents an important mechanism that controls global protein synthesis. Interestingly, our proteomics data show that proteins encoded by 5’TOP mRNAs are significantly repressed upon MEK1/2 inhibition in both HCT116 and SW480 cells (Fig. S4B).

Next, we sought to determine mRNA-specific changes in translation upon MEK1/2 inhibition, relative to the observed global translational repression. To this end, we performed Ribo-Seq in HCT116 cells treated with the MEK1/2 inhibitor for 24 h. After RNase I digest and purification of monosomes, ribosome protected fragments (ribosome footprints) were sequenced and displayed the characteristic three-nucleotide periodicity (Fig. S5A-B). Input RNA was randomly fragmented and sequenced in parallel to determine changes of mRNA abundance (Fig. S5C). Ribosome densities were calculated as the ratio of ribosome footprint to input RNA. In total,53 mRNAs showed a significant increase in ribosome density, whereas 169 showed a significant decrease. Among the translationally suppressed mRNAs, we identified 77 5’TOP mRNAs. While expression of these mRNAs is not reduced at the level of mRNA abundance, their reduced expression at the protein level as observed in our proteomics data is exclusively explained by translational downregulation (Fig. 5E, Fig. S5D). In contrast, we did not observe a significant translational regulation of AXIN1, other Wnt pathway components or Wnt-associated intestinal stem cell markers (*37*), of which many were transcriptionally induced upon MEK inhibition (yellow dots, Fig. 5F-G).

### mTOR inhibition decreases AXIN1 levels

Our aggregated results from the Ribo-Seq experiments (reduction of 5’TOP mRNAs) and immunoblot profiling (reduction in S6K and RPS6 phosphorylation) indicate that inhibition of the mTOR pathway plays a critical role in MEK1/2-induced translational repression. Hence, we hypothesized that mTOR inhibition may phenocopy the effect of MEK1/2 inhibitors on AXIN1 protein levels. To address this question, we treated HCT116 cells with the mTOR inhibitor Torin 1. Indeed, Torin 1 reduced AXIN1 protein levels in a dose-dependent manner, with 100 nM Torin 1 showing a similar effect as 100 nM MEK1/2 inhibitors (Fig. 6A). This result was confirmed using the clinically approved mTOR inhibitor rapamycin (Fig. 6B). Parallel measurements of RPS6 phosphorylation and the global translation rate by puromycin incorporation revealed that both MEK1/2 and mTOR inhibition led to a strong reduction of RPS6 phosphorylation and global protein biosynthesis (Fig. 6A-C). Since the reduction in AXIN1 levels was slightly stronger after MEK1/2 inhibition than after mTOR inhibition, we hypothesized that other pathways by which Ras-MAPK signaling controls translation may account for this phenotypic discrepancy. We therefore systematically combined mTOR inhibition with RSK1/2, S6K and MNK1/2 inhibitors and tested the effect of combination treatment on AXIN1 levels. We used the compounds BI-D1870 and LJI308 to inhibit RSK1/2, LY2584702 to target S6K1 and tomivosertib to inhibit MNK1/2. While the S6K1 inhibitor caused a minor reduction in RPS6 phosphorylation and AXIN1 levels, none of the other inhibitors enhanced the AXIN1 loss caused by mTOR inhibition (Fig. D-G). Finally, we measured changes in AXIN1 transcript levels upon mTOR inhibition. As opposed to MEK1/2 inhibitors, neither Torin-1 nor rapamycin repressed AXIN1 transcription at doses that reduced AXIN1 protein levels (Fig. 6H). From this, we concluded that loss of AXIN1 by mTOR inhibition is mediated by translational, not transcriptional repression.

**Figure 6:**
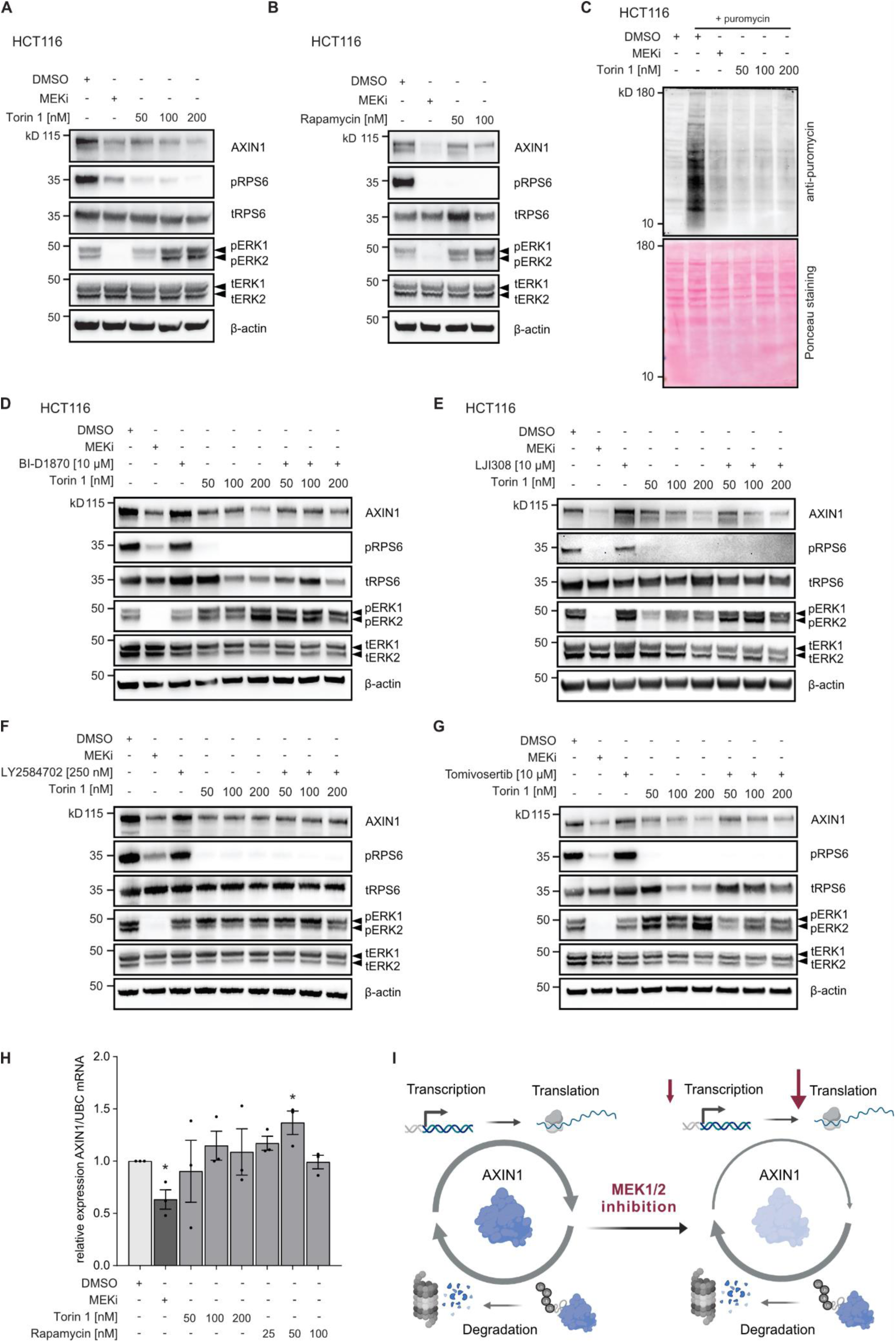
mTOR inhibition mimics effect of MEK inhibition on AXIN1 loss. **A-B,** The mTOR inhibitors Torin 1 and rapamycin dose-dependently decrease AXIN1 protein levels in the CRC cell line HCT116. Cells were treated with the indicated compounds for 24 h. **C**, The mTOR inhibitor Torin 1 decreases global translation rates similar to MEK1/2 inhibitors, as determined by puromycin incorporation assay. **D-G,** Effect of combinations of Torin 1 and inhibitors of central pathways by which Ras-MAPK signaling controls translation on AXIN1 levels. Torin 1 was combined with inhibitors of RSK (BI-D1870; LJI308), S6K1 (LY2584702) and MNK (tomivosertib). Effective concentrations of the inhibitors were selected based on literature. Only the S6K1 inhibitor induces a minor reduction of AXIN1 levels. No synergistic reduction of AXIN1 was observed for combinations of mTOR and other inhibitors. A-G: representative blots of three independent replicates are shown. **H**, mTOR inhibition does not reduce *AXIN1* mRNA levels. Data from three independent experiments are presented as mean ± SEM *p < 0.05, two-tailed Student’s t-test. Drug concentrations were as follows: MEKi - 100 nM trametinib. **I**, Model of MEK1/2-induced AXIN1 loss. Cellular AXIN1 levels are maintained by a dynamic balance of de novo synthesis and protein degradation. MEK1/2 inhibition reduces the transcription of AXIN1, and this loss is strongly enhanced by repression of global protein synthesis via an mTOR-dependent mechanism.

## DISCUSSION

AXIN1 is the central scaffold protein of the destruction complex, controls the integrity of this multi-protein complex and critically regulates Wnt signaling. Moreover, AXIN1 interacts with components of several other signaling pathways, including Hippo (*38*), AMPK (*39*) and TGF-beta (*40*), underlining its importance as a regulatory hub of cancer pathways. AXIN1 protein levels are tightly controlled by diverse cellular mechanisms, foremostly by posttranslational modifications and transcriptional regulation. We previously demonstrated that MEK1/2 inhibition activates Wnt signaling in CRC and proposed downregulation of AXIN1 as a potential underlying mechanism (*7*). A similar loss of AXIN1 was also observed upon BRAF and MEK inhibition in melanoma cells stimulated with Wnt3A (*41*, *42*), and in BRAF mutant CRC cell lines treated with BRAF inhibitors (*43*). However, how Ras-MAPK pathway inhibition mechanistically mediates AXIN1 loss is unknown. In the present study, we show that AXIN1 loss after MEK1/2 and GSK3B inhibition is commonly observed across CRC cell and organoid lines. On a mechanistic level, we found that GSK3B inhibition leads to AXIN1 protein degradation, while MEK1/2 inhibition did not affect protein stability of AXIN1. Instead, MEK1/2 inhibition results in global translational repression by an mTOR dependent mechanism. In line with this finding, mTOR inhibition reduced AXIN1 levels. Together, our results show that Ras-MAPK signaling critically maintains cellular AXIN1 protein homeostasis by translational control.

Due to the important role of AXIN1 as a regulatory hub of many cancer pathways, foremostly of Wnt signaling, levels of AXIN1 are extensively regulated by transcriptional and post-transcriptional mechanisms (*10*). Several transcription factors can activate transcription of *AXIN1* in a tissue-specific context, including GATA4 (*13*) and EGR1 (*16*). We previously showed that MEK1/2 inhibition represses *AXIN1* mRNA via the Ras-MAPK responsive transcription factor EGR1 in CRC cell lines, and that overexpression of EGR1 could prevent the effect of MEKi on *AXIN1* transcript levels (*7*). Our present results, however, suggest that additional mechanisms exist that further enhance AXIN1 loss upon MEK1/2 inhibition, as the observed reduction at the protein level was more pronounced than the transcriptional repression. AXIN1 protein homeostasis is well-known to be controlled by extensive post-translational modifications and subsequent degradation by the ubiquitin-proteasome system (*10*). More recently, the autophagy-lysosome pathway was implicated in AXIN1 protein turnover (*26*). Hence, we extensively investigated if active protein degradation is causative for the strong AXIN1 loss observed upon MEK1/2 inhibition. To this end, we performed cycloheximide chase assays, functional depletion of E3 ubiquitin ligases known to target AXIN1, and profiling of post-translational modifications of AXIN1 by ubiquitin-affinity precipitation. The aggregated results of these experiments indicate that, in contrast to GSK3B inhibition, AXIN1 loss by MEK1/2 inhibition is not dependent on ubiquitination-mediated protein degradation. This observation is further supported by our results demonstrating that inhibition of AXIN1 PARsylation by tankyrase inhibitors, which are known to stabilize AXIN1 proteins (*18*), prevents AXIN1 loss upon GSK3B inhibition more strongly than upon MEK1/2 inhibition. Therefore, we conclude that MEK1/2 inhibitors do not induce active protein degradation of AXIN1. Surprisingly, we found that pharmacological inhibition of lysosomal degradation using bafilomycin A1 also caused AXIN1 loss, indicating a more complex regulation of AXIN1 via the autophagy-lysosome pathway that needs further exploration.

To identify alternative mechanisms by which MEK1/2 inhibitors cause AXIN1 loss, we focused on regulation at the level of protein synthesis. The Ras-MAPK pathway is well known to control mRNA translation via MNK and RSK kinase family members (*44*). While MNKs cause eIF4E phosphorylation, eIF4B, eEF2K and RPS6 are phosphorylated upon activation of RSKs. Furthermore, the Ras-MAPK pathway can impinge on mTOR signaling to regulate translation, mainly via S6K, eIF4E-BP1 or LARP1 phosphorylation (*44*, *45*). This translation regulation is altered upon pharmacological perturbation of Ras-MAPK signaling. For instance, formation of the eukaryotic translation initiation complex eIF4F is decreased upon treatment with BRAF inhibitors in *BRAF* mutant melanoma, thyroid and CRC cells, whereas persistence of the complex is associated with resistance to MEK1/2 and BRAF inhibitors (*46*). Persistent formation of eIF4F can be mediated by re-activation of the Ras-MAPK pathway or continuous phosphorylation of eIF4E-BP1. Furthermore, suppression of mTOR, as evidenced by dephosphorylation of RPS6, was required for the apoptotic effect of BRAF and MEK1/2 inhibitors in *BRAF* mutant melanoma (*47*). These studies support a critical role of translational repression in mediating the antineoplastic effect of Ras-MAPK pathway inhibitors in cancer. Our results demonstrate that MEK1/2 inhibitors cause global translational repression in CRC, as shown by polysome profiling and puromycin incorporation assay. Mechanistically, we demonstrate that phosphorylation of mTOR targets (S6K, eIF4E-BP1) is reduced upon MEK1/2 inhibition. In line with this observation, our Ribo-Seq analysis revealed a strong translational repression of 5’TOP mRNAs, which are known to be regulated by mTOR signaling (*35*). In contrast, we did not observe an active translational regulation of genes associated with the Wnt signal pathway, including AXIN1. Finally, we show that treatment with mTOR inhibitors could phenocopy the effect of MEK1/2 inhibitors on AXIN1 levels. Thus, our results favor a model which proposes that AXIN1 protein homeostasis is maintained by continuous mRNA translation (Fig. 6I). Since we experimentally determined a protein half-life of 14 h for AXIN1, inhibition of translation by MEK1/2 inhibitors will result in a gradual reduction of AXIN1 levels within 24 h relative to the majority of proteins, which are much more stable (*24*), explaining the difference in kinetics of AXIN1 loss in comparison to GSK3B inhibitors.

Our observations show that the effects of Ras-MAPK on mRNA translation are mainly mediated by mTOR signaling in CRC. Direct interactions of Wnt and mTOR signaling have been described previously, with GSK3B being a critical link (*48*). Stimulation of the Wnt pathway by secreted ligands such as Wnt-1 can induce mTOR signaling in vitro and in vivo (*49*, *50*). This activation was caused by inhibition of GSK3B-mediated phosphorylation of TSC2 and was not dependent on beta-catenin (*50*). Conversely, interference with the mTOR pathway can also affect Wnt signaling. mTORC1 inhibition was shown to promote nuclear translocation of GSK3B, which was associated with a reduced function of the kinase in regulating MYC and Snail levels (*51*). mTORC1 was also found to negatively regulate ligand-stimulated Wnt/beta-catenin signaling by modulating the protein levels of FZD receptors. Pharmacological targeting of mTOR increased FZD2 expression, which was mediated by DVL (*52*). Here, we show that pharmacological targeting of mTOR results in AXIN1 loss in CRC. Given the widespread use of mTOR inhibitors in transplant medicine and cancer therapy, it is necessary to understand if our observations in cell and organoids models can be also confirmed in patient tissues.

Notably, GSK3B inhibition also induced a rapid loss of AXIN1, though via a mechanism distinct from MEK1/2 inhibition. GSK3B is known to phosphorylate AXIN1 at multiple sites, including T609 and S614 (*53*), and this phosphorylation is counteracted by PP2A (*54*, *55*). The functional consequences of GSK3B-mediated phosphorylation are diverse. In COS cells, overexpressed AXIN1 is destabilized upon treatment with the GSK3B inhibitor lithium (*53*). Mutation of two GSK3 phosphorylation sites (Ser322 and Ser326) to alanine or deletion of its GSK3B-binding site decreased the stability of AXIN1 in Xenopus egg extracts (*56*). Furthermore, modification of GSK3B phosphorylation sites reduced its interaction with beta-catenin and subsequent Wnt activation (*55*, *57*). Our results show that pharmacological GSK3B inhibition reduces AXIN1 levels in a large panel of CRC cell lines and in patient-derived organoids, indicating that previous observations with overexpressed AXIN1 in COS9 cells can be corroborated in cell lines with diverse genetic alterations of *APC* and *CTNNB1*. Mechanistically, GSK3B inhibition results in protein degradation of AXIN1, which was prevented by inhibition of PARsylation by tankyrase inhibitors. These results indicate that AXIN1 PARsylation might be a prerequisite for GSK3B-dependent regulation of AXIN1 phosphorylation and degradation, but the exact mechanisms will require further exploration.

In conclusion, our study highlights the importance of translational regulation in controlling the cellular levels of AXIN1, a central regulatory hub of several cancer pathways, whose turnover is subject to extensive regulation at multiple levels. Pharmacological perturbation of mTOR-dependent protein synthesis by MEK1/2 inhibitors can therefore reduce AXIN1 levels.

## MATERIAL AND METHODS

### Ethics approval

Experiments with patient-derived cancer organoids were approved by the Medical Ethics Committee II of the Medical Faculty Mannheim, Heidelberg University (Reference no. 2014– 633N-MA and 2016–607N-MA). Written informed consent was obtained from patients before performing any experimental procedures.

### Cell lines and culture

HCT116, SW480, SW837, HT29, HT55, COLO678, SNUC2A, and DLD1 cells were obtained from the American Type Culture Collection (ATCC). SW480, SW837, HT55, COLO678, SNUC2A, and DLD1 cells were cultured in RPMI 1640 medium (Gibco). HCT116 and HT29 cells were cultured in McCoy’s 5A medium (Gibco). All cell culture media were supplemented with 10% fetal bovine serum (FBS, Gibco), 1% Glutamax (Gibco), and 1% penicillin/streptomycin (Gibco). The absence of mycoplasma contamination was confirmed by regular PCR-based testing.

### Chemical compounds

Trametinib, CHIR99021, XAV939, MG132, bortezomib, bafilomycin A1, torin-1, rapamycin, LJI308, BI-D1870, LY2584702 and tomivosertib were all obtained from SelleckChem. Stock solutions for all drugs were prepared in DMSO and stored at −20°C.

### Organoid culture

Human organoid culture was performed as previously described (*7*). Human CRC organoids were embedded into BME (Trevigen) and maintained in Advanced DMEM/F12 medium (Gibco) supplemented with penicillin/streptomycin (Gibco), Glutamax (Gibco), and HEPES (= basal medium). The basal medium was further enriched with 100 ng/ml Noggin (PeproTech), 1 × B27 supplement (Life Technologies), 10 mM Nicotinamide (Sigma-Aldrich), 50 ng/ml human EGF (PeproTech), 10 nM Gastrin (PeproTech), 1 mM NAC (Sigma-Aldrich), 1 μM PGE2 (Tocris Bioscience), 500 nM A83-01 (Biocat), and 100 µg/ml Primocin (InvivoGen). The murine intestinal organoid line AKP (derived from C57B6 with Apc^fl/fl^, Kras^G12D/+^, Trp53^fl/fl^ mutational background) was previously published and kindly provided by Rene Jackstadt (*30*). Murine organoids were embedded into BME (Trevigen) and cultured in basal medium supplemented with 1% v/v N2 (Gibco), 2% v/v B27 (Gibco) and 100 ng/ml Noggin (PeproTech). Wild-type murine intestinal organoids were kindly provided by M. Schewe (Heidelberg University), embedded into BME and maintained in basal medium supplemented with 2.5% R-spondin conditioned medium, 2.5% Noggin, 2% N2, 0.2% Primocin, and 0.01% EGF. During thawing and passaging, 10 µM Y-27632 (SelleckChem) was included in the medium. Organoids were passaged every 4-7 d, and the culture medium was replenished every 2–3 d. Drug treatments were applied 72 h following seeding.

### Quantitative RT-PCR

RNA was extracted from cells or organoids using the peqGOLD Total RNA Kit (VWR Chemicals) following the manufacturer’s protocol. cDNA was synthesized using the Verso cDNA Synthesis Kit (Thermo Scientific) with 1 µg of total RNA as input. Quantitative PCR (qPCR) was performed using SYBR Green Master Mix (Applied Biosystems) on a StepOnePlus Real-Time PCR System (Applied Biosystems). Primers were designed using Primer3 and validated for efficiency and specificity.

### Western Blot

Cells were seeded at 5 × 10⁵ cells per well in 6-well plates and treated with the respective inhibitors. Protein lysates were prepared using saponin containing lysis buffer (PBS supplemented with 0.05% saponin (Sigma Aldrich), 2 mM EDTA (Sigma Aldrich), 10 mM β-mercaptoethanol) supplemented with proteinase inhibitors (Roche) and phosphatase inhibitor cocktail (1 and 2) (Sigma Aldrich). The protein concentration was determined by BCA assay (Thermo Fisher Scientific). Samples were boiled for 5 min at 99°C, proteins were separated by SDS-PAGE (BioRad) and transferred to a nitrocellulose membrane using the BioRad Turbo transfer system (BioRad). The membrane was blocked with 5% BSA in PBS-T (PBS supplemented with 0.1% Tween-20) for 1 hour, then incubated overnight at 4°C with primary antibodies, followed by secondary antibody incubation. Proteins were detected using SuperSignal chemiluminescent substrate (Thermo Fisher Scientific) and visualized using FUSION-SL-Advance imaging system (PeqLab). Membranes were stripped with Restore PLUS Western Blot Stripping Buffer (Thermo Fisher Scientific) and re-probed as needed.

### Cycloheximide chase assay

Cells were seeded at a density of 5 × 10⁵ cells per well in 6-well plates. After 24 h, cells were pretreated with inhibitors, followed by addition of 100 μg/ml cycloheximide (Sigma Aldrich) or DMSO. Cells were harvested at 0, 4, 8, and 12 h time points after treatment, washed with ice-cold PBS, and lysed in saponin-containing buffer (20 mM Tris-HCl pH 7.4, 130 mM NaCl, 2 mM EDTA, 10 mM β-mercaptoethanol, 0.05% saponin) supplemented with protease and phosphatase inhibitors. Lysates were processed for immunoblot as described above.

### Ubiquitin-affinity immunoprecipitation assay

A total of 2.5 × 10⁶ cells were seeded in 10 cm dishes and treated with the respective inhibitors, in the presence or absence of 10 µM MG132 (SelleckChem). Thereafter, cells were lysed using a lysis buffer containing 20 mM Tris–HCl, 130 mM NaCl, 10% glycerol, 2 mM EDTA, 1% Triton X-100, supplemented with protease and phosphatase inhibitors. The lysates were incubated with equilibrated agarose-TUBE2 beads (LifeSensors) overnight at 4°C. Control agarose beads (LifeSensors) were used in parallel to control for non-specific binding. After incubation, beads were collected by low-speed centrifugation at 3,000 × g and washed with TBS-T. Proteins were then eluted in SDS sample buffer. The eluates, along with input and unbound fractions, were analyzed by SDS-PAGE and immunoblot. For VU-101 antibody detection, membranes were pretreated with 0.5% glutaraldehyde (LifeSensors), blocked with 5% BSA, and incubated with the anti-ubiquitin antibody VU-1 (LifeSensors) overnight. After secondary antibody incubation, chemiluminescent detection was performed as described above.

### Co-Immunoprecipitation

A total of 2.5 × 10⁶ cells were seeded in 10 cm dishes. Twenty-four hours after seeding, cells were treated for 24 h with the respective inhibitors. After treatment, cells were washed with PBS and lysed using ice-cold lysis buffer containing 20 mM Tris–HCl, 130 mM NaCl, 10% glycerol, 2 mM EDTA, 1% Triton X-100, supplemented with protease and phosphatase inhibitors. Lysates were scraped, frozen overnight at -20°C, thawed, and clarified by centrifugation at 14,000 × g for 30 min at 4°C. 50 µl of Dynabeads Protein G magnetic beads (Thermo Fisher Scientific) were resuspended and incubated with 1 µg of primary antibody diluted in 200 µl of lysate and incubated overnight at 4°C with gentle rotation. After incubation, beads were separated using a magnet, and the supernatant was saved as the unbound fraction. The bead-antibody complex was washed six times with PBS containing 0.01% Tween-20 (Sigma Aldrich) to remove unspecific binding. Target antigens were eluted by heating the complex in 100 µl of 2x loading buffer at 99°C for 10 min, followed by magnetic separation. The supernatant was collected and analyzed by SDS-PAGE and immunoblot.

### RNA interference

HCT116 were seeded at a density of 5 × 10^5^ cells per well on 6-well plates. Twenty-four hours after cell seeding, cells were transfected with siGENOME SMARTPool non-targeting control siRNA#2 or siGENOME SMARTPool siRNAs against target genes (SIAH1, SIAH2, SMURF1, SMURF2, RNF146, TRIM65) (all from Horizon) and Lipofectamine RNAiMAX (Thermo Fisher Scientific) with a final concentration of 10 nM siRNA per well. For expression analysis, cells were harvested 48 h post transfection. For further treatment of transfected cells, the medium containing siRNAs was replaced 48 h after transfection by fresh medium, and drugs were added.

### Subcellular fractionation

HCT116 cells were seeded at a density of 5 × 10⁵ cells in 6-well plates and treated as indicated. Cells were then washed twice with ice-cold PBS and lysed in cytoplasmic extraction buffer (PBS supplemented with 0.05% saponin, 2 mM EDTA, 10 mM β-mercaptoethanol, protease inhibitors and freshly added phosphatase inhibitors). Lysates were incubated on ice for 15 min with gentle mixing and centrifuged at 16,000 × g at 4 °C for 15 min. The supernatant was collected as the cytoplasmic fraction. The remaining nuclear pellet was washed once with the same buffer, then resuspended in RIPA buffer (Thermo Fisher Scientific) with protease (Roche) and phosphatase inhibitors (Sigma Aldrich). The lysate was incubated for 30 min on ice with intermittent vortexing. Nuclear lysates were cleared by centrifugation at 16,000 × g for 15 min at 4 °C. Whole cell lysates were prepared in parallel using RIPA buffer. Fraction purity was validated by immunoblots using antibodies against β-actin (cytoplasmic marker) and histone H3 (nuclear marker).

### Mass Spectrometry

#### Preparation of cells

HCT116 cells were seeded in 10 cm culture dishes at a density of 2.5 × 10⁶ cells per dish. 24 h after seeding, cells were treated with 100 nM trametinib for 4 h. For GSK3B inhibition, cells were treated with 10 µM CHIR99021 for 30 min. Corresponding control cells were treated with DMSO for 30 min or 4 h. After treatment, cells were washed with cold PBS and lysed in ice-cold lysis buffer containing 20 mM Tris–HCl (pH 7.5), 130 mM NaCl, 10% glycerol, 2 mM EDTA, and 1% Triton X-100, supplemented with protease (Roche) and phosphatase (Sigma Aldrich) inhibitors. Lysates were scraped, frozen overnight at –20°C, thawed, and clarified by centrifugation at 14,000 × g for 30 minutes at 4°C. Pull-down of either AXIN1 or GSK3B was performed using Protein G Dynabeads (Thermo Fisher Scientific) conjugated with either AXIN1 or GSK3B antibody respectively, prepared in advance and kept at 4°C. For one immunoprecipitation reaction, 20 µl of Protein G Mag Sepharose conjugated with the corresponding antibody was added to the cell lysate and incubated overnight at 4°C. Next day, the tubes with the beads were briefly spinned down, placed on a magnetic rack and incubated for 5 min. The supernatant was discarded and the beads were washed three times with 1 ml ice-cold PBS buffer, pH 7.4 (Thermo Fisher Scientific). After the third wash, the immunoprecipitated protein complexes were stripped from the beads in two steps. In the first step, 25 µl of buffer containing 50 mM Tris pH 7.5; 1 M urea; 1 mM tris-(2-carboxyethyl)phosphine and 5 µg/ml of sequencing grade Trypsin (Promega) was added to the tubes with the beads and incubated for 30 min at 27 °C on a thermomixer at 900 rpm. The tubes were then placed on a magnetic rack and incubated for another 5 min at room temperature (RT). The supernatant was collected into fresh, labelled tubes. In the second step, the beads were washed twice with 25 µl of buffer containing 50 mM Tris pH 7.5; 1 M urea and 5 mM chloroacetamide and the supernatant was pooled with the supernatant from the step 1 in the corresponding tubes. The tubes were incubated overnight at 27°C to continue the digestion. Next day, the peptides generated by tryptic digestion were cleaned-up using SP3 method, as described elsewhere (*58*).

#### Preparation of antibody-conjugated Mag Sepharose beads

The appropriate volume of Protein G Mag Sepharose beads calculated for the number of IP reactions was incubated with either AXIN1 (C76H11) rabbit monoclonal antibody (Cat# 2087, Cell Signaling) or purified anti-GSK-3B mouse antibody (Cat# 610202, BD Biosciences). The concentration of antibody used for conjugation was 100 µg of antibody per 100 µl of settled beads. For the corresponding IP controls, the beads were conjugated with the isotypic IgG corresponding to the AXIN1 and GSK3B antibody, in particular with either rabbit IgG (Cat# 2729, Cell Signaling) or mouse (G3A1) IgG1 isotype control (Cat# 5415S, Cell Signaling). The IgG to bead ratio was identical, as described above.

#### Mass spectrometry analysis

The quantitative MS measurements were carried out using Dionex UltiMate 3000 UHPLC system (Thermo Fisher Scientific) coupled to an Exploris 480 Orbitrap mass spectrometer (Thermo Fisher Scientific). The peptides were separated by reverse-phase liquid chromatography with 0.1% formic acid (solvent A) and 100% acetonitrile supplemented with 0.1% formic acid (solvent B) as mobile phases, using a stepped gradient from 4% to 80% solvent B in 60 min on a nanoEasy M/Z peptide BEH C18 column (Waters, 250 mm × 75 μm 1/PK, 130 Å, 1.7 μm) mounted in the integrated column compartment of the UltiMate 3000 system heated to 55 °C. The peptides were eluted with a constant flow of 300 nl/min.

The Exploris 480 Orbitrap mass spectrometer was operated in DIA mode with a scan range of 350-1400 m/z, orbitrap resolution 120,000, normalized AGC target 300%, maxIT set to Auto mode and the precursors were analyzed in a sequence of 19 windows of variable width. The normalized HCD collision energy for the fragmentation of precursor ions was set to 28.

#### Data analysis of mass spectrometry

The files containing spectral data were analyzed using Spectronaut (version 19) software using a directDIA workflow against a nonredundant UniProt Human Proteome FASTA database from 30.01.2020 with the identification settings as follows: precursor Q-value cutoff 0.01, precursor posterior error probability (PEP) cutoff 0.2, protein Q-value cutoff (experiment-wise) 0.01, protein Q-value cutoff (run-wise) 0.05, protein PEP cutoff 0.75. Carbamidomethylation was set as a fixed modification, and acetylation (N-term), oxidation, phosphorylation, and ubiquitination as variable modifications. For the quantification, the data were normalized based on a retention time-dependent local regression model as previously described (*59*), with precursor filtering based on identified Q-value, and maxLFQ quantification method based on interrun peptide ratios. The proteins were grouped by protein group ID and peptides were grouped by a stripped peptide sequence of the identified precursors. The missing values were imputed with 0.001 in the resulting datasets using a python script.

### CRISPR-mediated gene knockout

A previously validated sgRNA sequence targeting RNF146 (5’-TGAGCGCACTAGTAGAGAGC-3’) was selected to generate gene knockouts (*28*). Oligonucleotides encoding the sgRNA were synthesised by Eurofins Inc. Oligonucleotides were phosphorylated, annealed and cloned into px459 plasmid (#62988, Addgene) using a published protocol (24157548). To generate RNF146 knockouts, HCT116 cells were seeded in 6-well plates and transiently transfected with 1 µg of px459 encoding sgRNF146 per well using Lipofectamine 3000 (Thermo Fischer Scientific). Seventy-two hours post-transfection, cells were selected with 1 µg/ml of puromycin (Gibco) for 72 h. Puromycin was then removed and cells were allowed to grow until formation of colonies. Surviving colonies were pooled and knockout of RNF146 was validated by Sanger sequencing.

### Polysome profiling

HCT116 cells were seeded at a density of 3 ×10^6^ per dish in 10 cm dishes 24 h before treatment with the respective inhibitors. Before lysis, cells were incubated with 100 μg/ml cycloheximide (CHX) for 5 min at RT and washed with ice-cold PBS before harvesting by scraping in polysome lysis buffer (20 mM Tris-HCl pH 7.4, 5 mM MgCl2, 150 mM NaCl, 1 mM DTT, 1 % Triton-X-100, 200 U/ml RNasin [Promega], EDTA-free complete protease inhibitors [Roche]). The lysates were rotated for 10 min at 4°C and cleared from cell debris by centrifugation at 10,000 × g for 10 min at 4°C. For Western Blot analysis, 40 µl of lysate per condition was set aside. The remaining lysate (250 µl) was loaded onto linear 17.5–50% [w/v] sucrose gradients (dissolved in 20 mM Tris-HCl pH 7.5, 5mM MgCl2 and 150 mM NaCl) and centrifuged in an SW60 rotor (Beckman) for 2 h at 40,000 rpm and 4°C. Polysome profiles were recorded by detection of UV absorbance at 254 nm using an Äkta Prime system in conjunction with an siFractor (siTOOLs Biotech). Profiles were aligned along the 80S peak, normalized and quantified using the QuAPPro application (*60*). The ratio of polysomal ribosomes was calculated by dividing the area under the polysomal part of the curve by the total area under the curve starting from the 40S peak.

### Ribosome profiling

HCT116 cells were seeded at a density of 3 × 10^6^ per dish in 10 cm dishes 24 h before treatment with 100 nM trametinib (MEKi) or DMSO. Before lysis, cells were incubated with 100 μg/ml cycloheximide (CHX) for 5 min at RT and washed with ice-cold PBS before harvesting by scraping in polysome lysis buffer (without RNasin). The lysates were rotated for 10 min at 4°C and cleared from cell debris by centrifugation at 10,000 × g for 10 min at 4°C. Per condition, 30 µl of lysate were set aside for Western Blot analysis, and another 30 µl for isolation of input RNA. Lysates were digested with 240 U RNase I (Ambion) per A260 Unit for 20 min at 4°C. After 17.5–50% [w/v] sucrose density gradient centrifugation, fractions of ∼300 µl were collected during gradient elution and supplemented with 300 µl urea buffer (10 mM Tris, pH 7.5, 350 mM NaCl, 10 mM EDTA, 1% SDS, and 7 M urea) and 300 µl phenol:chloroform:isoamyl alcohol (25:24:1). By phase separation, RNA of the cytoplasmic lysates (inputs) and the monosomal fractions (ribosome protected fragments; RPF) was isolated. The input and RPF samples were depleted of ribosomal RNA (rRNA) using the Human-Mouse-Rat Ribo-Seq riboPOOL kit (siTOOLs Biotech). After random fragmentation of input RNA at 95°C for 12 min in alkaline fragmentation buffer (2 mM EDTA, 88 mM NaHCO3, 12 mM Na2CO3), both input and footprint samples were size-selected (25-35 nt) on 15% polyacrylamide Tris-borate-EDTA-urea gels. Following end-repair using T4 PNK, 1.65 ng per sample were used for library preparation using the NEBNext Multiplex Small RNA Library Prep Set according to the manufacturer’s instructions. Libraries were multiplexed and sequenced as 80 nt long single-end reads on a NextSeq 550 sequencing device (Illumina). After removal of adapter sequences (AGATCGGAAGAGCACACGTCTGAACTCCAGTCAC) with the FASTX-toolkit (http://hannonlab.cshl.edu/fastx_toolkit/), the sequences were aligned to human rRNA and tRNA sequences using Bowtie v1.2.2 (*61*), allowing a maximum of two mismatches per read and reporting all alignments in the best stratum. Reads that did not align in this step were mapped to the basic set of Gencode V38. Read counts were summarized at the gene level by counting all 25–35 nt long reads that align to annotated ORFs of one specific gene (as defined by a common gene symbol). Relative to the 5′ end of the read, an offset of -12 nt to the start codon and -15 nt for the stop codon was assumed (Fig. S5B). Differentially translated mRNAs were identified with a likelihood ratio test followed by multiple testing correction with the Benjamini-Hochberg procedure in DESeq2 v1.46.0 (*62*).

### RNA sequencing

RNA was isolated from cultured cells using the peqGOLD Total RNA Kit (VWR), following the manufacturer’s instructions. After lysis in TRK buffer and homogenization, lysates were applied to RNA Mini Columns. DNase I digestion was performed to remove genomic DNA. RNA was washed, eluted with preheated nuclease-free water (70 °C) and stored at −70 °C. Libraries were prepared at the DKFZ Genomics & Proteomics Core Facility using the TruSeq Stranded mRNA Library Prep Kit (Illumina) and sequenced on a NovaSeq 6000 (Illumina) as paired-end 100 nt long reads. Alignment to hg38 was performed using STAR v2.5.3a (*63*) allowing a maximum of 10 mismatches and providing a gtf with exon coordinates of the Gencode Basic V38 transcript set. Alignments were annotated and summarized at the gene level using featureCounts within the subread package v1.6.3 (*64*). Differential expression was tested by pair-wise comparison of each MEKi time-point to its respective DMSO control using the DESeq2 package v1.46.0. Principal component analyses were performed using the vst() and plotPCA() functions of the DESeq2 package (*65*).

### Puromycin incorporation assay

Cells were seeded at a density of 5 × 10⁵ in 6-well plates. Twenty-four hours after seeding, cells were treated for 24 h with the indicated drugs. Ten minutes before the end of treatment, 5 µg/ml puromycin (Gibco) was added for 10 min at 37°C. Cells were then washed twice with PBS, and resuspended in lysis buffer. For immunoblot analysis, equal amounts of total cell lysates were separated by SDS-PAGE. Puromycin signals were detected with an anti-puromycin antibody. The signal intensity was measured along the entire lane and normalized to the Ponceau S staining of the corresponding lane.

## ACKNOWLEDGEMENT

We thank the NGS Core Facility of the German Cancer Research Center and the NGS core facility at the Institute for Clinical Chemistry at the Medical Faculty Mannheim at Heidelberg University for help with RNA sequencing. We thank Rene Jackstadt (German Cancer Research Center) and Matthias Schewe (Medical Faculty Mannheim, Heidelberg University) for providing intestinal organoid models.

## AUTHOR CONTRIBUTIONS

Conceptualization: JS, TZ. Methodology: NV, LW, JS, TZ. Investigation: NV, LW, NM, RI, JK, OS, AL, YC, JB, KB, JS. Visualization: NV, LW, YC, JS, TZ. Funding acquisition: TZ. Project administration: JS, TZ. Supervision: JS, TZ. Writing - original draft: JS, TZ. Writing - review & editing: NV, LW, NM, RI, JK, OS, AL, YC, JB, KB, JS, MB, GS, MPE, JS, TZ

## DATA AND MATERIAL AVAILABILITY

Data and material will be made available upon request.

## REFERENCES

1. R. L. Siegel, N. S. Wagle, A. Cercek, R. A. Smith, A. Jemal, Colorectal cancer statistics, 2023. CA Cancer J Clin 73, 233–254 (2023).

2. M. Riihimaki, A. Hemminki, J. Sundquist, K. Hemminki, Patterns of metastasis in colon and rectal cancer. Sci Rep 6, 1–9 (2016).

3. A. Cervantes, R. Adam, S. Roselló, D. Arnold, N. Normanno, J. Taïeb, J. Seligmann, T. De Baere, P. Osterlund, T. Yoshino, E. Martinelli, Metastatic colorectal cancer: ESMO Clinical Practice Guideline for diagnosis, treatment and follow-up ☆. Annals of Oncology 34, 10–32 (2023).

4. S. Kopetz, J. Desai, E. Chan, J. R. Hecht, P. J. O’Dwyer, D. Maru, V. Morris, F. Janku, A. Dasari, W. Chung, J. P. J. Issa, P. Gibbs, B. James, G. Powis, K. B. Nolop, S. Bhattacharya, L. Saltz, Phase II pilot study of vemurafenib in patients with metastatic BRAF-mutated colorectal cancer. Journal of Clinical Oncology 33, 4032–4038 (2015).

5. J. R. Infante, L. A. Fecher, G. S. Falchook, S. Nallapareddy, M. S. Gordon, C. Becerra, D. J. DeMarini, D. S. Cox, Y. Xu, S. R. Morris, V. G. R. Peddareddigari, N. T. Le, L. Hart, J. C. Bendell, G. Eckhardt, R. Kurzrock, K. Flaherty, H. A. Burris, W. A. Messersmith, Safety, pharmacokinetic, pharmacodynamic, and efficacy data for the oral MEK inhibitor trametinib: a phase 1 dose-escalation trial. Lancet Oncol 13, 773–81 (2012).

6. M. B. Ryan, R. B. Corcoran, Therapeutic strategies to target RAS-mutant cancers. Nat Rev Clin Oncol 15, 709–720 (2018).

7. T. Zhan, G. Ambrosi, A. M. Wandmacher, B. Rauscher, J. Betge, N. Rindtorff, R. S. Häussler, I. Hinsenkamp, L. Bamberg, B. Hessling, K. Müller-Decker, G. Erdmann, E. Burgermeister, M. P. Ebert, M. Boutros, MEK inhibitors activate Wnt signalling and induce stem cell plasticity in colorectal cancer. Nat Commun 10, 2197 (2019).

8. T. Zhan, N. Rindtorff, M. Boutros, Wnt signaling in cancer. Oncogene 36, 1461–1473 (2017).

9. V. S. W. Li, S. S. Ng, P. J. Boersema, T. Y. Low, W. R. Karthaus, J. P. Gerlach, S. Mohammed, A. J. R. Heck, M. M. Maurice, T. Mahmoudi, H. Clevers, Wnt Signaling through Inhibition of β-Catenin Degradation in an Intact Axin1 Complex. Cell 149, 1245–1256 (2012).

10. L. Qiu, Y. Sun, H. Ning, G. Chen, W. Zhao, Y. Gao, The scaffold protein AXIN1: gene ontology, signal network, and physiological function. Cell Commun Signal 22 (2024).

11. H. K. Arnold, X. Zhang, C. J. Daniel, D. Tibbitts, J. Escamilla-Powers, A. Farrell, S. Tokarz, C. Morgan, R. C. Sears, The Axin1 scaffold protein promotes formation of a degradation complex for c-Myc. EMBO J 28, 500–512 (2009).

12. W. Liu, H. Rui, J. Wang, S. Lin, Y. He, M. Chen, Q. Li, Z. Ye, S. Zhang, C. C. Siu, Y. G. Chen, J. Han, S. C. Lin, Axin is a scaffold protein in TGF-β signaling that promotes degradation of Smad7 by Arkadia. EMBO Journal 25, 1646–1658 (2006).

13. S. Suthon, R. S. Perkins, J. Lin, J. R. Crockarell, G. A. Miranda-Carboni, S. A. Krum, GATA4 and estrogen receptor alpha bind at SNPs rs9921222 and rs10794639 to regulate AXIN1 expression in osteoblasts. Hum Genet 141, 1849–1861 (2022).

14. N. O. Chimge, G. H. Little, S. K. Baniwal, H. Adisetiyo, Y. Xie, T. Zhang, A. O’Laughlin, Z. Y. Liu, P. Ulrich, A. Martin, P. Mhawech-Fauceglia, M. J. Ellis, D. Tripathy, S. Groshen, C. Liang, Z. Li, D. E. Schones, B. Frenkel, RUNX1 prevents oestrogen-mediated AXIN1 suppression and β-catenin activation in ER-positive breast cancer. Nat Commun 7 (2016).

15. D. N. Jackson, K. M. Alula, Y. Delgado-Deida, R. Tabti, K. Turner, X. Wang, K. Venuprasad, R. F. Souza, L. Desaubry, A. L. Theiss, The synthetic small molecule FL3 combats intestinal tumorigenesis via axin1-mediated inhibition of Wnt/b-catenin signaling. Cancer Res 80, 3519–3529 (2020).

16. M. Zhang, Y. Liao, B. Lönnerdal, EGR-1 is an active transcription factor in TGF-β2-mediated small intestinal cell differentiation. Journal of Nutritional Biochemistry 37, 101–108 (2016).

17. Y. Zhang, S. Liu, C. Mickanin, Y. Feng, O. Charlat, G. A. Michaud, M. Schirle, X. Shi, M. Hild, A. Bauer, V. E. Myer, P. M. Finan, J. A. Porter, S. M. A. Huang, F. Cong, RNF146 is a poly(ADP-ribose)-directed E3 ligase that regulates axin degradation and Wnt signalling. Nat Cell Biol 13, 623–629 (2011).

18. S.-M. A. Huang, Y. M. Mishina, S. Liu, A. Cheung, F. Stegmeier, G. A. Michaud, O. Charlat, E. Wiellette, Y. Zhang, S. Wiessner, M. Hild, X. Shi, C. J. Wilson, C. Mickanin, V. Myer, A. Fazal, R. Tomlinson, F. Serluca, W. Shao, H. Cheng, M. Shultz, C. Rau, M. Schirle, J. Schlegl, S. Ghidelli, S. Fawell, C. Lu, D. Curtis, M. W. Kirschner, C. Lengauer, P. M. Finan, J. A. Tallarico, T. Bouwmeester, J. A. Porter, A. Bauer, F. Cong, Tankyrase inhibition stabilizes axin and antagonizes Wnt signalling. Nature 461, 614–20 (2009).

19. S. Kim, E. H. Jho, The protein stability of Axin, a negative regulator of Wnt signaling, is regulated by Smad ubiquitination regulatory factor 2 (Smurf2). Journal of Biological Chemistry 285, 36420–36426 (2010).

20. L. Ji, B. Jiang, X. Jiang, O. Charlat, A. Chen, C. Mickanin, A. Bauer, W. Xu, X. Yan, F. Cong, The SIAH E3 ubiquitin ligases promote Wnt/β-catenin signaling through mediating Wnt-induced Axin degradation. Genes Dev 31, 904–915 (2017).

21. Y. F. Yang, M. F. Zhang, Q. H. Tian, C. Z. Zhang, TRIM65 triggers β-catenin signaling via ubiquitylation of Axin1 to promote hepatocellular carcinoma. J Cell Sci 130, 3108–3115 (2017).

22. T. T. H. Lui, C. Lacroix, S. M. Ahmed, S. J. Goldenberg, C. A. Leach, A. M. Daulat, S. Angers, The Ubiquitin-Specific Protease USP34 Regulates Axin Stability and Wnt/β-Catenin Signaling. Mol Cell Biol 31, 2053–2065 (2011).

23. L. Ji, B. Lu, R. Zamponi, O. Charlat, R. Aversa, Z. Yang, F. Sigoillot, X. Zhu, T. Hu, J. S. Reece-Hoyes, C. Russ, G. Michaud, J. S. Tchorz, X. Jiang, F. Cong, USP7 inhibits Wnt/β-catenin signaling through promoting stabilization of Axin. Nat Commun 10 (2019).

24. J. Li, Z. Cai, L. P. Vaites, N. Shen, D. C. Mitchell, E. L. Huttlin, J. A. Paulo, B. L. Harry, S. P. Gygi, Proteome-wide mapping of short-lived proteins in human cells. Mol Cell 81, 4722–4735.e5 (2021).

25. Y. Haupt, R. Maya, A. Kazaz, M. Oren, Mdm2 promotes the rapid degradation of p53. Nature 387, 296–299 (1997).

26. M. V. Gammons, M. Renko, J. E. Flack, J. Mieszczanek, M. Bienz, Feedback control of wnt signaling based on ultrastable histidine cluster co-aggregation between Naked/NKD and axin. Elife 9, 1–22 (2020).

27. M. G. Callow, H. Tran, L. Phu, T. Lau, J. Lee, W. N. Sandoval, P. S. Liu, S. Bheddah, J. Tao, J. R. Lill, J. A. Hongo, D. Davis, D. S. Kirkpatrick, P. Polakis, M. Costa, Ubiquitin ligase RNF146 regulates tankyrase and Axin to promote Wnt signaling. PLoS One 6 (2011).

28. L. Nie, C. Wang, N. Li, X. Feng, N. Lee, D. Su, M. Tang, F. Yao, J. Chen, Proteome-wide Analysis Reveals Substrates of E3 Ligase RNF146 Targeted for Degradation. Molecular and Cellular Proteomics 19 (2020).

29. K. Imkeller, G. Ambrosi, N. Klemm, A. Claveras Cabezudo, L. Henkel, W. Huber, M. Boutros, Metabolic balance in colorectal cancer is maintained by optimal Wnt signaling levels. Mol Syst Biol 18 (2022).

30. N. Vaquero-Siguero, N. Schleussner, J. Volk, M. Mastel, J. Meier, R. Jackstadt, Modeling Colorectal Cancer Progression Reveals Niche-Dependent Clonal Selection. Cancers (Basel) 14 (2022).

31. Q. Xiao, J. E. Riedesser, T. Mulholland, Z. Li, J. Buchloh, P. Albrecht, M. Li, N. Venkatachalam, O. Skabkina, A. Klupsch, E. Eichhorn, L. Wang, S. Belle, N. Schulte, D. Schmitz, M. F. Froelich, E. Valentini, K. E. Boonekamp, Y. Petersen, T. Miersch, E. Burgermeister, C. Herskind, M. R. Veldwijk, C. Brochhausen, R. Ihnatko, J. Krijgsveld, I. Kurth, M. Boutros, M. P. Ebert, T. Zhan, J. Betge, An organoid platform reveals MEK-PARP co-targeting to enhance radiation response in rectal cancer. bioRxiv, 2024.06.06.597640 (2024).

32. H. Lavoie, J. Gagnon, M. Therrien, ERK signalling: a master regulator of cell behaviour, life and fate. Nat Rev Mol Cell Biol 21, 607–632 (2020).

33. S. Shin, L. Wolgamott, J. Tcherkezian, S. Vallabhapurapu, Y. Yu, P. P. Roux, S. O. Yoon, Glycogen synthase kinase-3β positively regulates protein synthesis and cell proliferation through the regulation of translation initiation factor 4E-binding protein 1. Oncogene 33, 1690–1699 (2014).

34. S. Shin, L. Wolgamott, Y. Yu, J. Blenis, S. O. Yoon, Glycogen synthase kinase (GSK)-3 promotes p70 ribosomal protein S6 kinase (p70S6K) activity and cell proliferation. Proc Natl Acad Sci U S A 108 (2011).

35. B. D. Fonseca, C. Zakaria, J. J. Jia, T. E. Graber, Y. Svitkin, S. Tahmasebi, D. Healy, H. D. Hoang, J. M. Jensen, I. T. Diao, A. Lussier, C. Dajadian, N. Padmanabhan, W. Wang, E. Matta-Camacho, J. Hearnden, E. M. Smith, Y. Tsukumo, A. Yanagiya, M. Morita, E. Petroulakis, J. L. González, G. Hernández, T. Alain, C. K. Damgaard, La-related protein 1 (LARP1) represses terminal oligopyrimidine (TOP) mRNA translation downstream of mTOR complex 1 (mTORC1). Journal of Biological Chemistry 290, 15996–16020 (2015).

36. O. Meyuhas, T. Kahan, The race to decipher the top secrets of TOP mRNAs. Biochim Biophys Acta Gene Regul Mech 1849, 801–811 (2015).

37. A. Merlos-Suárez, F. M. Barriga, P. Jung, M. Iglesias, M. V. Céspedes, D. Rossell, M. Sevillano, X. Hernando-Momblona, V. Da Silva-Diz, P. Muñoz, H. Clevers, E. Sancho, R. Mangues, E. Batlle, The intestinal stem cell signature identifies colorectal cancer stem cells and predicts disease relapse. Cell Stem Cell 8, 511–524 (2011).

38. E. J. Lee, E. Seo, J. W. Kim, S. A. Nam, J. Y. Lee, J. Jun, S. Oh, M. Park, E. H. Jho, K. H. Yoo, J. H. Park, Y. K. Kim, TAZ/Wnt-β-catenin/c-MYC axis regulates cystogenesis in polycystic kidney disease. Proc Natl Acad Sci U S A 117, 29001–29012 (2020).

39. Y. L. Zhang, H. Guo, C. S. Zhang, S. Y. Lin, Z. Yin, Y. Peng, H. Luo, Y. Shi, G. Lian, C. Zhang, M. Li, Z. Ye, J. Ye, J. Han, P. Li, J. W. Wu, S. C. Lin, AMP as a low-energy charge signal autonomously initiates assembly of axin-ampk-lkb1 complex for AMPK activation. Cell Metab 18, 546–555 (2013).

40. R. Xie, D. Yi, D. Zeng, Q. Jie, Q. Kang, Z. Zhang, Z. Zhang, G. Xiao, L. Chen, L. Tong, D. Chen, Specific deletion of Axin1 leads to activation of β-catenin/BMP signaling resulting in fibular hemimelia phenotype in mice. Elife 11 (2022).

41. T. L. Biechele, R. M. Kulikauskas, R. A. Toroni, O. M. Lucero, R. D. Swift, R. G. James, N. C. Robin, D. W. Dawson, R. T. Moon, A. J. Chien, Wnt/β-catenin signaling and AXIN1 regulate apoptosis triggered by inhibition of the mutant kinase BRAFV600Ein human melanoma. Sci Signal 5, ra3–ra3 (2012).

42. W. H. Conrad, R. D. Swift, T. L. Biechele, R. M. Kulikauskas, R. T. Moon, A. J. Chien, Regulating the response to targeted MEK inhibition in melanoma: enhancing apoptosis in NRAS- and BRAF-mutant melanoma cells with Wnt/β-catenin activation. Cell Cycle 11, 3724–30 (2012).

43. G. Chen, C. Gao, X. Gao, D. H. Zhang, S.-F. Kuan, T. F. Burns, J. Hu, Wnt/β-catenin pathway activation mediates adaptive resistance to BRAF inhibition in colorectal cancer. Mol Cancer Ther 17, molcanther.0561.2017 (2017).

44. P. P. Roux, I. Topisirovic, Signaling Pathways Involved in the Regulation of mRNA Translation. Mol Cell Biol 38 (2018).

45. C. G. Proud, Phosphorylation and signal transduction pathways in translational control. Cold Spring Harb Perspect Biol 11 (2019).

46. L. Boussemart, H. Malka-Mahieu, I. Girault, D. Allard, O. Hemmingsson, G. Tomasic, M. Thomas, C. Basmadjian, N. Ribeiro, F. Thuaud, C. Mateus, E. Routier, N. Kamsu-Kom, S. Agoussi, A. M. Eggermont, L. Désaubry, C. Robert, S. Vagner, eIF4F is a nexus of resistance to anti-BRAF and anti-MEK cancer therapies. Nature 513, 105–109 (2014).

47. R. B. Corcoran, S. M. Rothenberg, A. N. Hata, A. C. Faber, A. Piris, R. M. Nazarian, R. D. Brown, J. T. Godfrey, D. Winokur, J. Walsh, M. Mino-Kenudson, S. Maheswaran, J. Settleman, J. A. Wargo, K. T. Flaherty, D. A. Haber, J. A. Engelman, TORC1 suppression predicts responsiveness to RAF and MEK inhibition in BRAF-mutant melanoma. Sci Transl Med 5 (2013).

48. A. Prossomariti, G. Piazzi, C. Alquati, L. Ricciardiello, Are Wnt/β-Catenin and PI3K/AKT/mTORC1 Distinct Pathways in Colorectal Cancer? CMGH 10, 491–506 (2020).

49. R. M. Castilho, C. H. Squarize, L. A. Chodosh, B. O. Williams, J. S. Gutkind, mTOR Mediates Wnt-Induced Epidermal Stem Cell Exhaustion and Aging. Cell Stem Cell 5, 279–289 (2009).

50. K. Inoki, H. Ouyang, T. Zhu, C. Lindvall, Y. Wang, X. Zhang, Q. Yang, C. Bennett, Y. Harada, K. Stankunas, C. yu Wang, X. He, O. A. MacDougald, M. You, B. O. Williams, K. L. Guan, TSC2 Integrates Wnt and Energy Signals via a Coordinated Phosphorylation by AMPK and GSK3 to Regulate Cell Growth. Cell 126, 955–968 (2006).

51. S. J. Bautista, I. Boras, A. Vissa, N. Mecica, C. M. Yip, P. K. Kim, C. N. Antonescu, mTOR complex 1 controls the nuclear localization and function of glycogen synthase kinase 3β. Journal of Biological Chemistry 293, 14723–14739 (2018).

52. H. Zeng, B. Lu, R. Zamponi, Z. Yang, K. Wetzel, J. Loureiro, S. Mohammadi, M. Beibel, S. Bergling, J. Reece-Hoyes, C. Russ, G. Roma, J. S. Tchorz, P. Capodieci, F. Cong, MTORC1 signaling suppresses Wnt/β-catenin signaling through DVL-dependent regulation of Wnt receptor FZD level. Proc Natl Acad Sci U S A 115, E10362–E10369 (2018).

53. H. Yamamoto, S. Kishida, M. Kishida, S. Ikeda, S. Takada, A. Kikuchi, Phosphorylation of axin, a Wnt signal negative regulator, by glycogen synthase kinase-3β regulates its stability. Journal of Biological Chemistry 274, 10681–10684 (1999).

54. E. T. Strovel, D. Wu, D. J. Sussman, Protein phosphatase 2Cα dephosphorylates axin and activates LEF-1-dependent transcription. Journal of Biological Chemistry 275, 2399–2403 (2000).

55. K. Willert, S. Shibamoto, R. Nusse, Wnt-induced dephosphorylation of Axin releases β-catenin from the Axin complex. Genes Dev 13, 1768–1773 (1999).

56. M. J. Cantoria, E. Alizadeh, J. Ravi, R. P. Varghese, N. Bunnag, K. W. Pond, A. N. Kettenbach, Y. Ahmed, A. L. Paek, J. J. Tyson, K. Doubrovinski, E. Lee, C. A. Thorne, Feedback in the β-catenin destruction complex imparts bistability and cellular memory. Proc Natl Acad Sci U S A 120 (2023).

57. E. H. Jho, S. Lomvardas, F. Costantini, A GSK3β phosphorylation site in Axin modulates interaction with β-catenin and Tcf-mediated gene expression. Biochem Biophys Res Commun 266, 28–35 (1999).

58. C. S. Hughes, S. Moggridge, T. Müller, P. H. Sorensen, G. B. Morin, J. Krijgsveld, Single-pot, solid-phase-enhanced sample preparation for proteomics experiments. Nat Protoc 14, 68–85 (2019).

59. S. J. Callister, M. A. Dominguez, C. D. Nicora, X. Zeng, C. L. Tavano, S. Kaplan, T. J. Donohue, R. D. Smith, M. S. Lipton, Application of the accurate mass and time tag approach to the proteome analysis of sub-cellular fractions obtained from Rhodobacter sphaeroides 2.4.1. aerobic and photosynthetic cell cultures. J Proteome Res 5, 1940–1947 (2006).

60. C. Schiller, S. Reitter, J. A. Lehmann, K. Fenzl, J. Schott, QuAPPro: An R/shiny app for Quantification and Alignment of Polysome Profiles. bioRxiv, 2024.05.02.592260 (2024).

61. B. Langmead, C. Trapnell, M. Pop, S. L. Salzberg, Ultrafast and memory-efficient alignment of short DNA sequences to the human genome. Genome Biol 10 (2009).

62. M. I. Love, W. Huber, S. Anders, Moderated estimation of fold change and dispersion for RNA-seq data with DESeq2. Genome Biol 15 (2014).

63. A. Dobin, C. A. Davis, F. Schlesinger, J. Drenkow, C. Zaleski, S. Jha, P. Batut, M. Chaisson, T. R. Gingeras, STAR: Ultrafast universal RNA-seq aligner. Bioinformatics 29, 15–21 (2013).

64. M. Kearse, R. Moir, A. Wilson, S. Stones-Havas, M. Cheung, S. Sturrock, S. Buxton, A. Cooper, S. Markowitz, C. Duran, T. Thierer, B. Ashton, P. Meintjes, A. Drummond, Geneious Basic: An integrated and extendable desktop software platform for the organization and analysis of sequence data. Bioinformatics 28, 1647–1649 (2012).

65. M. I. Love, W. Huber, S. Anders, Moderated estimation of fold change and dispersion for RNA-seq data with DESeq2. Genome Biol 15 (2014).

