## SUPPLEMENTARY MATERIALS for "MEK inhibition induces AXIN1 loss in colorectal cancer by mTOR associated suppression of protein synthesis"

#### Supplementary Figures

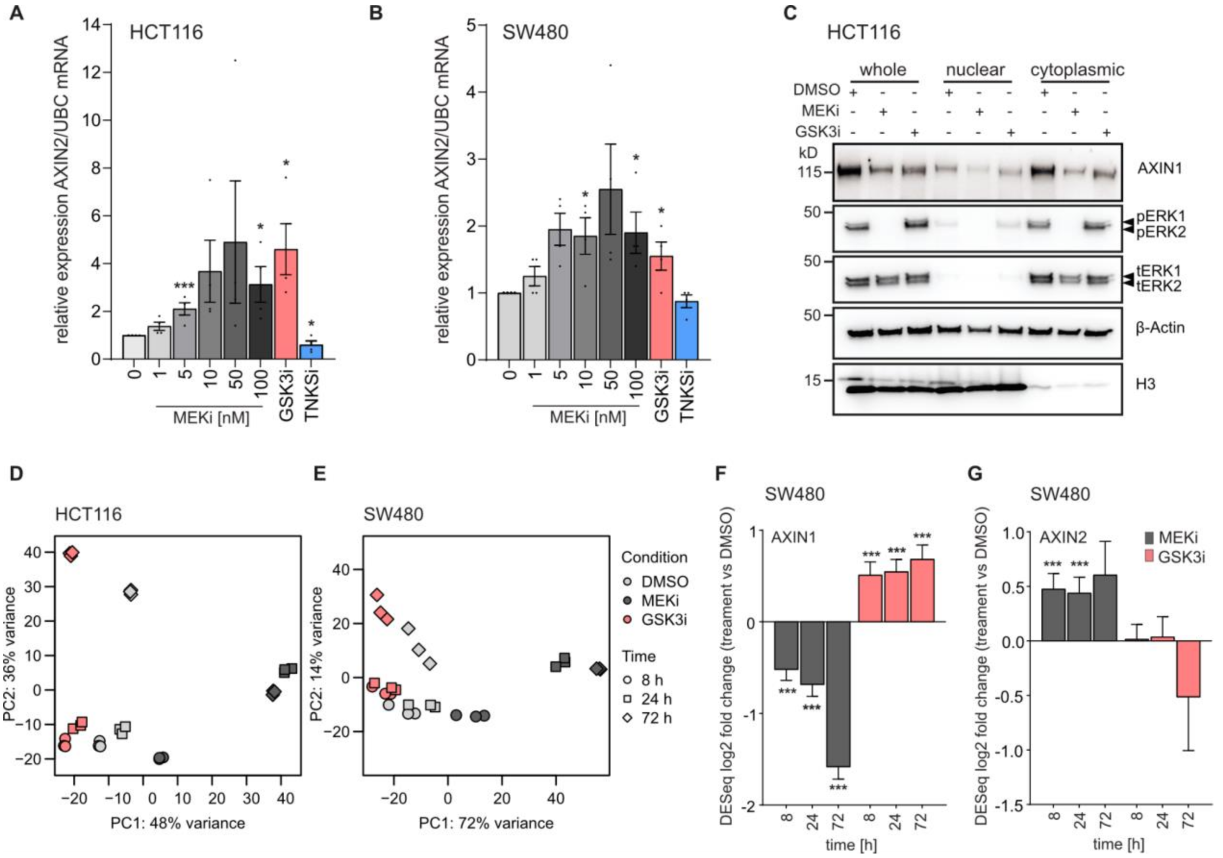


**Figure S1: Effect of MEK and GSK3 inhibition on expression of Wnt pathway-associated genes. A,** Concentration-dependent effect of MEK1/2 inhibition on *AXIN2* mRNA levels in HCT116 after treatment for 24 h. **B,** Concentration-dependent effect of MEK1/2 inhibition on *AXIN2* mRNA levels in SW480 after treatment for 24 h. A-B: Data from three independent experiments are presented as mean ± SEM *p < 0.05, **p < 0.01, ***p < 0.001, two-tailed Student's t-test. **C**, Subcellular fractionation reveals loss of AXIN1 protein upon MEKi in both nuclear and cytoplasmic fractions. Cells were treated for 24 h with MEKi before subcellular fractionation. **D-E**, PCA plots of RNAseq results of HCT116 (D) and SW480 (E) treated with GSK3i and MEKi for the indicated time periods. **F-G**, Time-dependent effect of MEK1/2 and GSK3B inhibition on *AXIN1* (F) and *AXIN2* (G) mRNA levels in SW480, analyzed by RNAseq. Cells were treated for the indicated time periods with the inhibitors. Statistical analysis was performed using DESeq2. Data from three independent experiments are presented as mean ± SD ***p < 0.001. Drug concentrations were as follows: MEKi - 100 nM trametinib, GSK3Bi - 10 µM CHIR-99021.

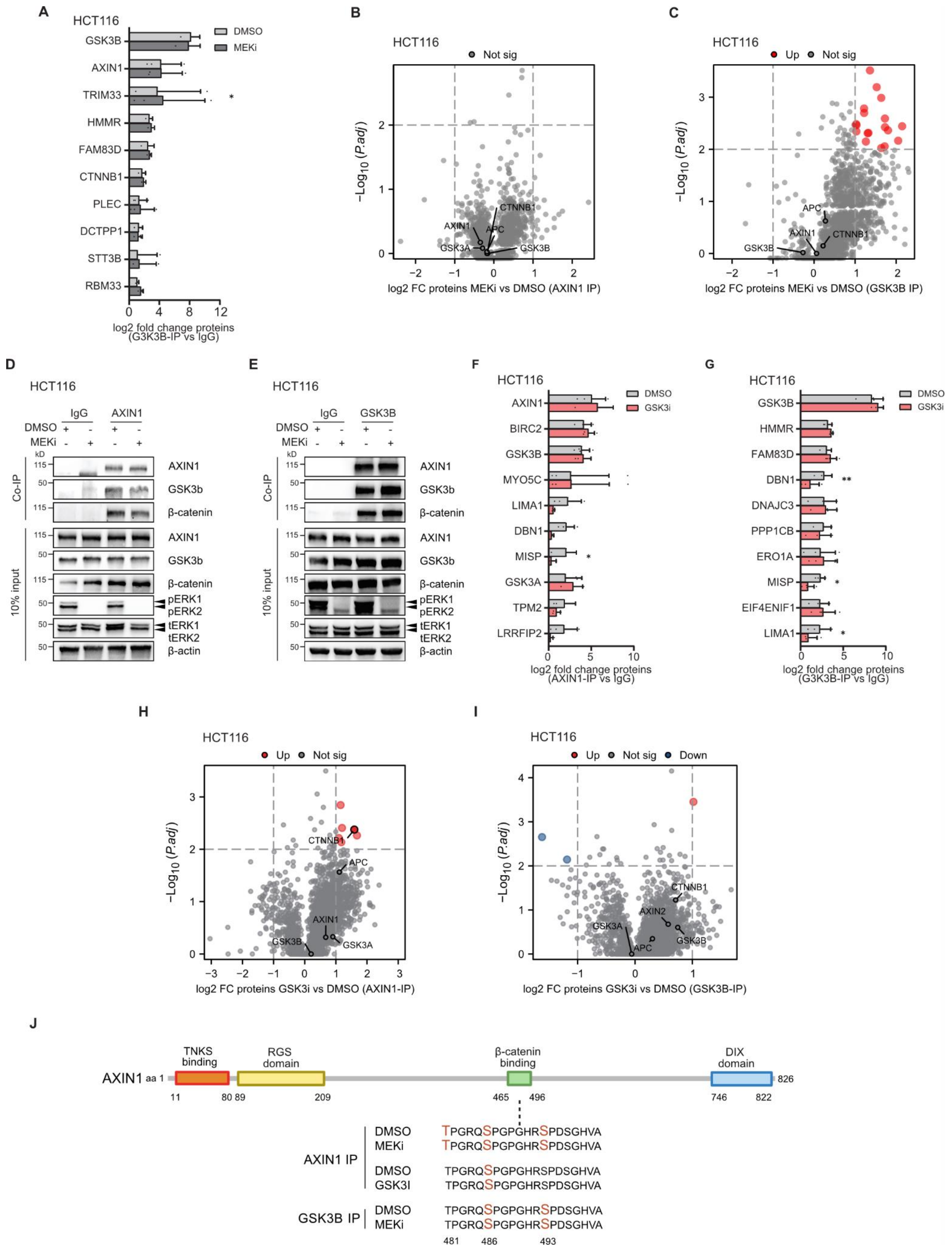

**Figure S2: Protein interactions and posttranslational modifications of AXIN1 and GSK3B after MEK1/2 and GSK3B inhibition.**

**A,** No changes in protein-protein interactions of GSK3B with main destruction complex components after MEKi. HCT116 were treated for 4 h with 100 nM trametinib (MEKi), followed by Co-IP with anti-GSK3B antibody or IgG control and mass spectrometry. Ten most enriched proteins after Co-IP with anti-GSK3B antibody are shown. **B-C**, Volcano plots showing differential protein-protein interactions after Co-IP with anti-AXIN1 (B) and anti-GSK3B (C) antibodies following MEKi. Destruction complex members are highlighted with names. **D-E**, No change of interactions between AXIN1, GSK3B and

CTNNB1 upon short time treatment with MEKi. HCT116 were treated for 4 h with 100 nM trametinib and affinity purification of lysates was performed using antibodies against AXIN1 (D) and GSK3B (E). Representative images of three independent replicates are shown. **F-G**, Changes in protein-protein interactions of AXIN1 and GSKBi with specific binding partners after GSK3i. HCT116 were treated for 30 min with 10 µM CHIR-99021 (GSK3i), followed by Co-IP with anti-AXIN1 (F), anti-GSK3B antibodies (G) or IgG control and mass spectrometry. Ten most enriched proteins after Co-IP with anti-AXIN1 and anti-GSK3B antibodies are shown. **H-I**, Volcano plots showing differential protein-protein interactions after IP with anti-AXIN1 (H) and anti-GSK3B (I) antibodies following GSK3i. **J**, Detected phosphorylation sites in AXIN1 after MEKi and GSK3i by mass spectrometry. Only sites with a probability of 0.99 and occurring in at least two of three (MEKi) or four (GSK3i) replicates are shown. Mass spectrometry data from n = 3 (MEKi) and n = 4 (GSK3i) independent replicates are presented as mean ± SD $*p < 0.05$, $**p < 0.01$, two-tailed Student's t-test.

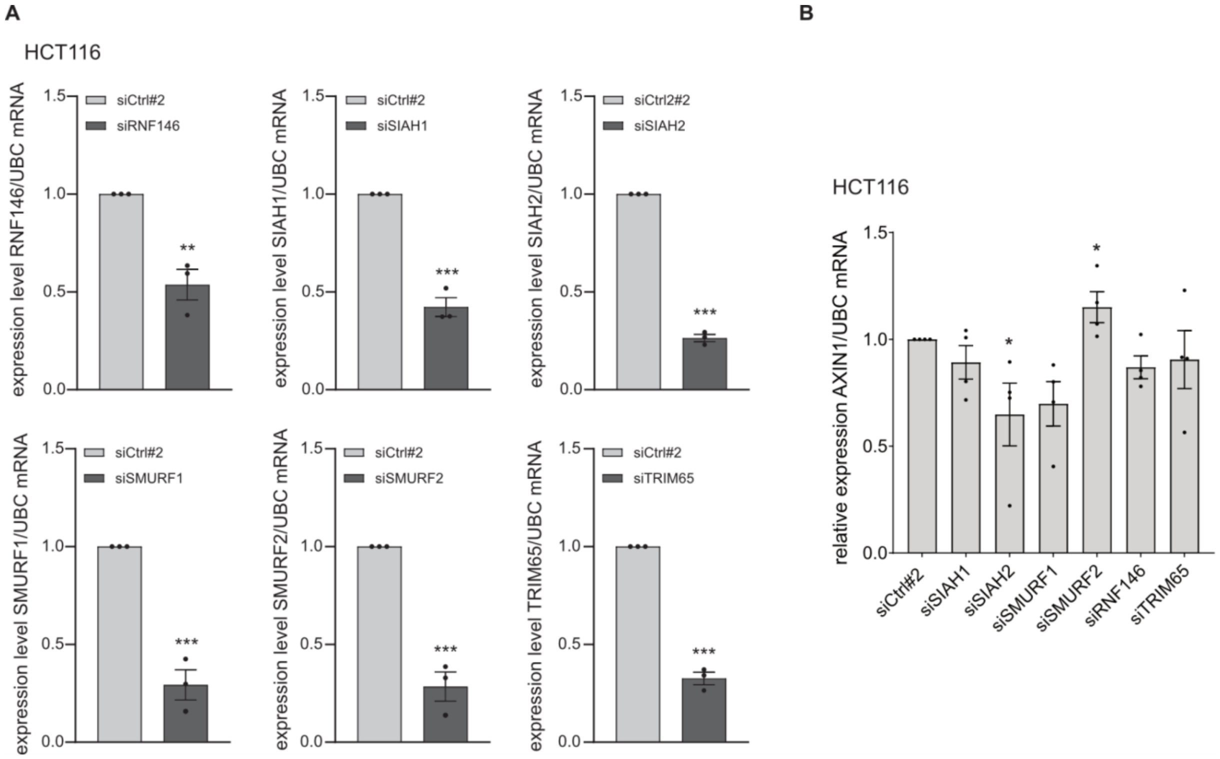


**Figure S3: Efficiency of RNAi-mediated knockdown of E3 ubiquitin ligases.**

**A,** Knockdown efficiency of siRNAs targeting AXIN1-associated E3 ubiquitin ligases. Expression of target genes were determined 48 h post transfection of pooled siRNAs. Non-targeting siRNAs (siCtrl#2) were used as control. **B,** Effect of knockdown of AXIN1-associated E3 ubiquitin ligases on expression of AXIN1. AXIN1 transcript levels were determined 48 h post transfection of pooled siRNAs. A-B, Data from 3-4 independent experiments are presented as mean ± SEM $*p < 0.05$, two-tailed Student's t-test.

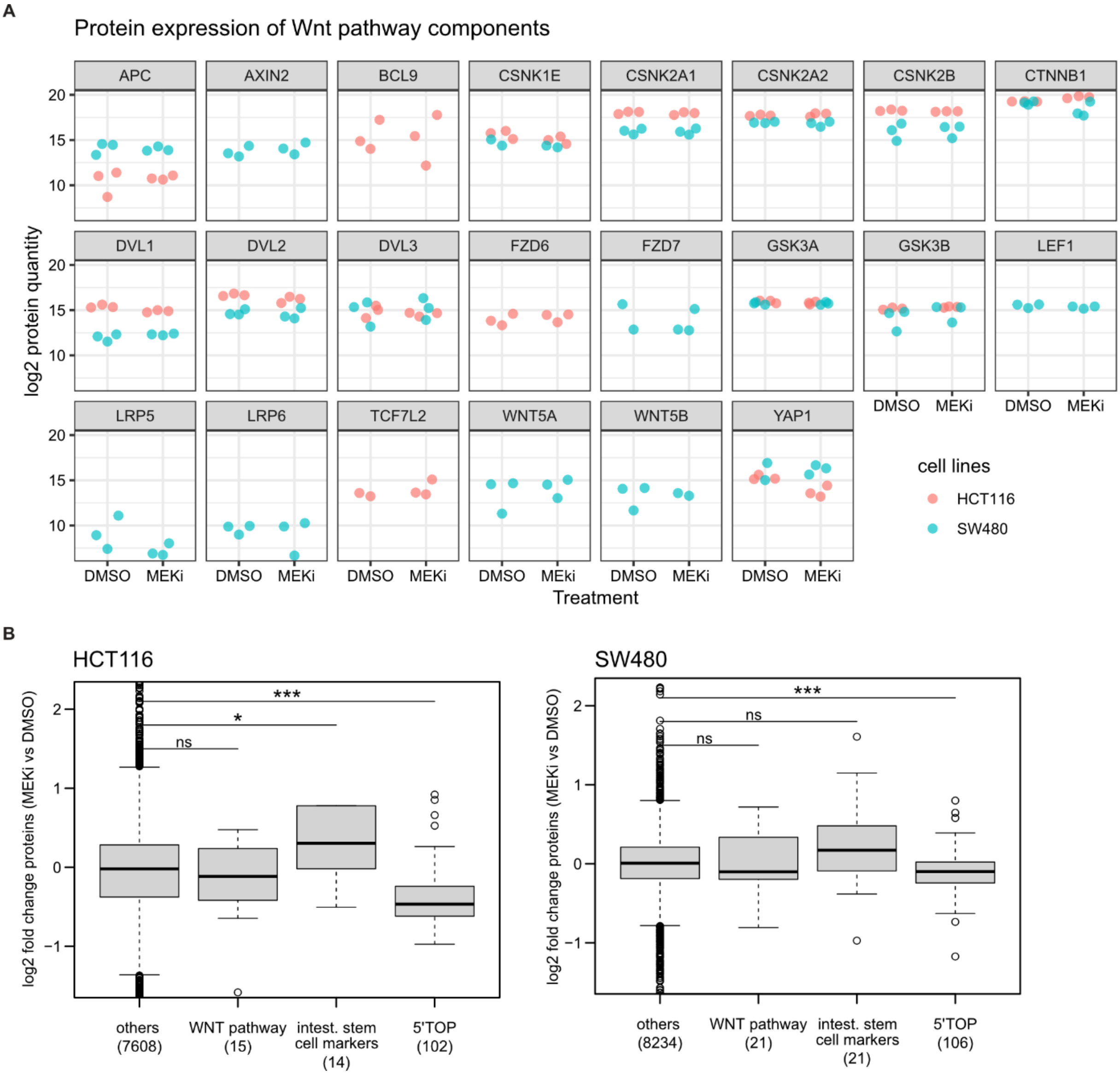


**Figure S4: Protein levels of Wnt pathway components upon MEK1/2 inhibition.**

**A,** Protein abundance of selected Wnt pathway components in HCT116 and SW480 cells treated with 100 nM trametinib (MEKi) for 24 h, detected by global proteome profiling. Three biological replicates were analyzed for each cell line. Only Wnt pathway components that are detected in at least two of three replicates of each cell line are shown. No significant changes of protein abundances between DMSO and MEKi were detected. The dataset is derived from Xiao et. al (*31*). **B,** Fold-changes of protein abundances derived from the same dataset as in (A) for Wnt pathway components, Wnt-associated intestinal stemness markers and proteins encoded by 5'TOP mRNAs. Differences were tested using a two-sided Wilcoxon rank sum test (*p < 0.05, ***p < 0.001).

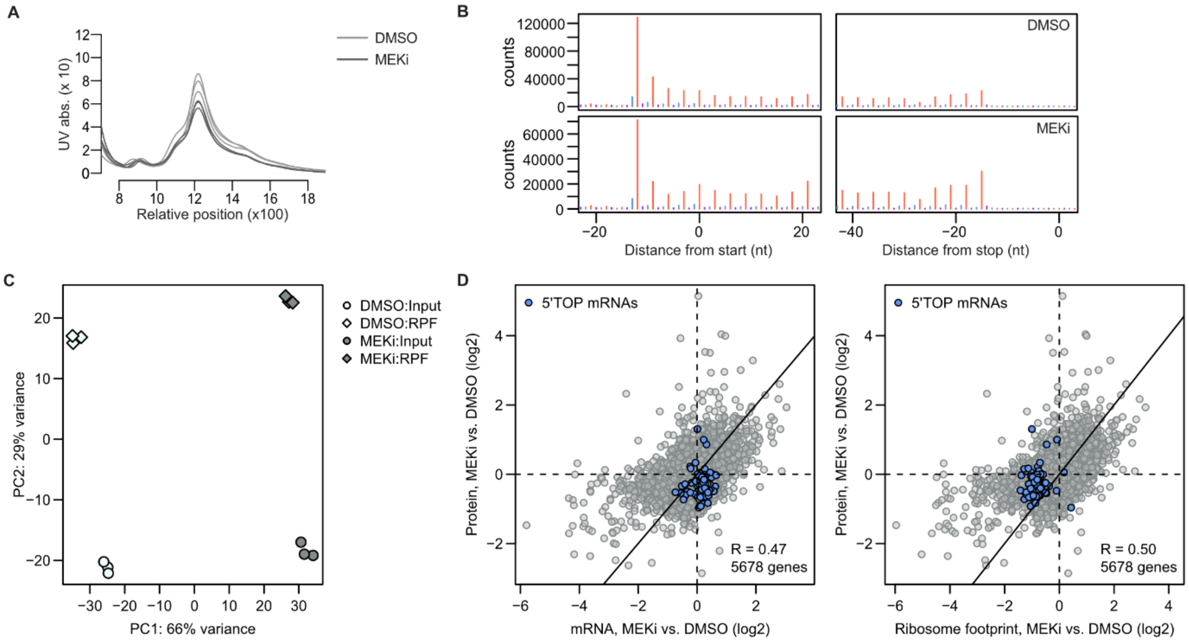


**Figure S5: Quality control of Ribo-Seq experiments.**

**A,** UV absorption profiles recorded after RNase I digest and sucrose-density gradient ultracentrifugation for isolation of monosomes. **B,** Number of read counts at the indicated distances from the start and stop codon relative to the 5' end of the ribosome protected fragment for 30 nt long reads. The annotated reading frame is highlighted in red. **C,** Principal component analysis of the normalized read counts. **D,** Relationship between protein fold-changes and mRNA (left panel) or ribosome footprint fold-changes (right panel), as determined from the mean of three biological replicates. R: Pearson's correlation coefficient.

### Supplementary Tables

#### Supplementary Table 1: List of antibodies for immunoblot and immunoprecipitation

| Antibody | Company | Catalogue # | Species | Dilution |
|---|---|---|---|---|
| Axin1 (C76H11) | Cell Signaling Technology | 2087 | rabbit | 1:2000 |
| phospho-p44/42 MAPK (Erk1/2) (Thr202/Tyr204) | Cell Signaling Technology | 4370 | rabbit | 1:2000 |
| p44/42 MAPK (Erk1/2) (Thr202/Tyr204) | Cell Signaling Technology | 9102 | rabbit | 1:2000 |
| β-actin (C4) HRP | Santa Cruz Biotechnology | sc-47778 HRP | mouse | 1:20000 |
| GSK-3β | BD Biosciences | 610202 | mouse | 1:2000 |
| beta-catenin | BD Biosciences | 610154 | mouse | 1:2000 |
| histone H3 (D1H2) HRP | Cell Signaling Technology | 12648S | rabbit | 1:2000 |
| p53 | Cell Signaling Technology | 9282S | rabbit | 1:2000 |
| LC3B | Cell Signaling Technology | 2775S | rabbit | 1:2000 |
| phospho-eIF4E (Ser209) | Cell Signaling Technology | 9741 | rabbit | 1:2000 |
| eIF4E | Cell Signaling Technology | 9742 | rabbit | 1:2000 |
| eIF4E-BP1 (53H11) | Cell Signaling Technology | 9644 | rabbit | 1:2000 |
| phospho-eIF4E-BP1 (Thr37/46) | Cell Signaling Technology | 2855 | rabbit | 1:2000 |
| S6 kinase | Cell Signaling Technology | 2708 | rabbit | 1:1000 |
| phospho-S6 Kinase | Cell Signaling Technology | 9206 | mouse | 1:1000 |
| RPS6 | Cell Signaling Technology | 2217 | rabbit | 1:1000 |
| phospho-RPS6 | Cell Signaling Technology | 4858 | rabbit | 1:2000 |
| tubulin | Sigma-Aldrich | T9026 | mouse | 1:5000 |
| □-actin | Abcam | ab8227 | rabbit | 1:4000 |

| | | | | |
|---|---|---|---|---|
| anti-rabbit IgG, HRP linked | Cell Signaling Technology | 7074 | goat | 1:5000 |
| anti-mouse IgG, HRP linked | Cell Signaling Technology | 7076 | horse | 1:5000 |
| anti-ubiquitin (VU-1) | LifeSensors | VU-0101 | mouse | 1:2000 |
| anti-tankyrase-1/2 (E-10) | Santa Cruz Biotechnology | sc-365897 | mouse | 1:2000 |
| anti-puromycin, clone 12D10 | Sigma-Aldrich | MABE343 | mouse | 1:5000 |

**Supplementary Table 2: List of primers for quantitative PCR**

| Target gene | Species | Forward primer | Reverse primer |
|---|---|---|---|
| AXIN1 | human | ATGGAGCTCTCCGAGACAGA | TAGTACGCCACAACGATGCT |
| AXIN2 | human | AGTGTGAGGTCCACGGAAAC | CTGGTGCAAAGACATAGCCA |
| UBC | human | CTGATCAGCAGAGGTTGATCTTT | TCTGGATGTTGTAGTCAGACAGG |
| TRIM65 | human | CGCCAACCGTCACTTCTATCT | ACAGGGTCAGGGTCCTACC |
| SMURF1 | human | ATTCGATAACCATTAGCGTGTGG | CGCCGGTTCCTATTCTGTCTC |
| SMURF2 | human | GGCAATGCCATTCTACAGATACT | CAACCGAGAAATCCAGCACCT |
| SIAH1 | human | TGTTTGTAGCAACTGTCGCC | AGCCACTTTCTCCATAGCCA |
| SIAH2 | human | GCCATCGTCCTGCTCATTGGCA | ACCAATATGGGAAGGCAGGCAGGAAGGGGC |

| | | | |
|---|---|---|---|
| RNF146 | human | ATTCCCGAGGATTTCCTTGACA | GCTCATCGTACTGCCACCA |
| TNKS1 | human | TGGTGCTGATGTTCATGCAAA | ACAAGCTCCATGCTTTAGTAGC |
| TNKS2 | human | GTGAATGCCCAAGACAAAGGAGG | GGTGTGAAAGCCCATTTGTCCG |
| SDHA | mouse | TGTTCAGTTCCACCCCACA | TCTCCACGACACCCTTCTGT |
| AXIN2 | mouse | AGGATGCTGAAGGCTCAAAG | TCGCCTTCTTGAAATAATACCTG |
| LGR5 | mouse | CTTCACTCGGTGCAGTGCT | GATCAGCCAGCTACCAAATAGG |
| ASCL2 | mouse | GAGAGCTAAGCCCGATGGA | AGGTCCACCAGGAGTCACC |
